# Predicting the functional impact of single nucleotide variants in *Drosophila melanogaster* with FlyCADD

**DOI:** 10.1101/2025.02.27.640642

**Authors:** Julia Beets, Julia Höglund, Bernard Y. Kim, Jacintha Ellers, Katja M. Hoedjes, Mirte Bosse

## Abstract

Understanding how genetic variants drive phenotypic differences is a major challenge in molecular biology. Single nucleotide polymorphisms form the vast majority of genetic variation and play critical roles in complex, polygenic phenotypes, yet their functional impact is poorly understood from traditional gene-level analyses. In-depth knowledge about the impact of single nucleotide polymorphisms has broad applications in health and disease, population genomic and evolution studies. The wealth of genomic data and available functional genetic tools make *Drosophila melanogaster* an ideal model species for studies at single nucleotide resolution. However, to leverage these resources for genotype-phenotype research and potentially combine it with the power of functional genetics, it is essential to develop techniques to predict functional impact and causality of single nucleotide variants.

Here, we present FlyCADD, a functional impact prediction tool for single nucleotide variants in *D. melanogaster*. FlyCADD, based on the Combined Annotation-Dependent Depletion (CADD) framework, integrates over 650 genomic features - including conservation scores, GC content, and DNA secondary structure - into a single metric reflecting a variant’s predicted impact on evolutionary fitness. FlyCADD provides impact prediction scores for any single nucleotide variant on the *D. melanogaster* genome. We demonstrate the power of FlyCADD for typical applications, such as the ranking of phenotype-associated variants to prioritize variants for follow-up studies, evaluation of naturally occurring polymorphisms, and refining of CRISPR-Cas9 experimental design. FlyCADD provides a powerful framework for interpreting the functional impact of any single nucleotide variant in *D. melanogaster*, thereby improving our understanding of genotype-phenotype connections.

**Article summary:** Single nucleotide polymorphisms (SNPs), the most common form of genomic variation, drive micro-evolution and adaptation. In *Drosophila melanogaster*, many SNPs are associated with phenotypes, yet functional validation is rare and experimentally challenging. FlyCADD is a new impact prediction tool that integrates *D. melanogaster* genome annotations into a single score predicting SNP impact. FlyCADD can be applied to distinguish causal from neutral variants, prioritize variants prior to functional studies, and to interpret natural variation, thereby improving understanding of genotype-phenotype relationships.

## Introduction

Unravelling the causative relationships between phenotypes, genes and their variants represents a significant challenge in molecular biology, especially at the resolution of single nucleotides (Cano-Gamez and Trynka 2020; Gallagher and Chen-Plotkin 2018). Species typically harbor millions of single nucleotide polymorphisms (SNPs) in their genomes; for instance, over five million SNPs have been identified in the fruit fly, *Drosophila melanogaster* (Auton et al. 2015; Nunez et al. 2024; Wang et al. 2015). Genomic studies, such as genome-wide association studies (GWAS) and Evolve and Resequence (E&R) studies, have identified vast numbers of SNPs associated with traits of interest. This led to the discovery of many candidate loci associated to a broad range of phenotypes, for example Type 2 diabetes and loci linked to fitness strategies such as behavioral choices in insect pupation (Uffelmann et al. 2021; Visscher et al. 2017; Wangler et al. 2017; Xue et al. 2018; Zhang et al. 2020). Elucidating the functional effects of individual single nucleotide variants is critical for understanding how genetic variation shapes phenotypes and advances our understanding of evolutionary adaptation, population dynamics, disease susceptibility, and other fundamental biological processes (de Visser and Krug 2014; Ungerer et al. 2008). Yet, despite the unprecedented resolution and availability of genotype data, the functional impact of individual SNPs is only rarely elucidated after identification of phenotype-associated variants (Hoedjes et al. 2023; Hoedjes et al. 2022; Katzenberger et al. 2015; Perlmutter et al. 2024; Schaid et al. 2018; Visscher et al. 2017).

There are several technical and biological challenges that complicate interpretation of genetic variation at single nucleotide level. Many of the SNPs identified in association studies are genetically linked variants that are not under selection themselves but merely show allele frequency (AF) changes by hitchhiking with loci that are under selection (Cano-Gamez and Trynka 2020; Gallagher and Chen-Plotkin 2018; Smith and Haigh 1974). Epistatic interactions between loci, environmental factors and additive effects of SNPs can further complicate the interpretation of the effect(s) of individual SNPs (Gallagher and Chen-Plotkin 2018; Huang et al. 2012; Yashin et al. 2012). Statistical identification of SNPs associated with traits can lead to false positives if non-causal SNPs appear significant due to linkage. False negatives can occur if for example true causal variants are masked by complex genetic architecture. As a result, statistical associations do not always reflect the context in which variants occur and, therefore, cannot reliably be taken as an indication for a functional relationship between a variant and a phenotype. For understanding any genotype-phenotype link it is imperative to distinguish causal SNPs from (linked) neutral loci, yet distinguishing these two types of variants among phenotype-associated SNPs remains a significant challenge (Franssen et al. 2015; Smith and Haigh 1974; Wang et al. 2022). Developing techniques and tools to assess the functional impact of SNPs is therefore essential for advancing genotype-phenotype research at SNP-level.

Functional genetic tests such as gene knockdown or knockout, through RNAi or CRISPR-Cas9 for example are typically applied to study the function of genetic loci of interest (Akhund-Zade et al. 2017; Hoedjes et al. 2023; Hoedjes et al. 2022; Mokashi et al. 2021; Parker et al. 2020; Wolf et al. 2023). These techniques are often applied at the gene-level, meaning that they focus on (nearby) genes associated with the SNP of interest, but do not directly test the candidate SNP(s) itself (Hoedjes et al. 2023; Hoedjes et al. 2022; Mokashi et al. 2021; Perlmutter et al. 2024). Such gene-level approaches might not identify effects of the targeted SNPs but rather all phenotypic effects that a gene might have (Hoedjes et al. 2023; Hoedjes et al. 2022). As different SNPs within the same gene could potentially have different functional impact or pleiotropic effects, it is important to understand the functional impact of SNPs instead of genes to understand phenotypic variation within species (Hoedjes et al. 2023; Mokashi et al. 2021; Zhang and Yang 2015). Precise genome editing, for example using CRISPR-Cas9, enables targeted investigations at single nucleotide resolution, but these methods are costly and time-consuming so only few SNPs can be tested. This is one of the reasons why experimental validation of candidate SNPs remains scarce even though functional validation provides the most direct evidence for SNP function (de Visser and Krug 2014; Hoedjes et al. 2023; Mokashi et al. 2021; Perlmutter et al. 2024). To be able to focus functional validation on the most promising SNPs, without spending costly time and resources on SNPs without functional impact, it is critical to narrow down the set of candidates based on expected functional impact.

Computational approaches can be instrumental in prioritization of candidate SNPs prior to functional studies. Existing computational approaches for predicting the effects of SNPs include SIFT (Ng and Henikoff 2001), PolyPhen-2 (Adzhubei et al. 2010), VEP (McLaren et al. 2016), SnpEff (Cingolani et al. 2012), ANNOVAR (Wang et al. 2010), AlphaMissense (Cheng et al. 2023), ProteoCast (Abakarova et al. 2025) and GPN-MSA (Benegas et al. 2023). These tools are each based on a different narrow set of genomic annotations—primarily conservation scores, genomic structure, and locus proximity. Moreover, they differ at the scale at which they are applicable, focusing on variants in the whole genome, (nearby) genes, or coding sequence (Wang et al. 2022). Studies on SNP impact typically integrate insights from multiple tools to take advantage of various annotations (Gibert et al. 2017; Hall et al. 2019). Yet, many additional annotations, such as regulatory elements, amino acid changes, chromatin states, and DNA secondary structures, remain underutilized, and could provide essential insights for predicting the functional significance of genetic variants at single-nucleotide resolution (de Visser and Krug 2014; Öztürk-Çolak et al. 2024).

A promising computational approach that combines a broad range of annotations to predict functional impact of genetic variants is the Combined Annotation-Dependent Depletion (CADD) framework (Kircher et al. 2014). The CADD framework is a first of its kind tool integrating diverse annotations into a single metric reflecting predicted functional impact of any SNP across the genome. CADD differs from other VEPs like SIFT (Ng and Henikoff 2001) or REVEL (Wang et al. 2022), which have been developed based on pathogenic variants in (mostly human) coding sequence. Although these VEPs outperform CADD on amino acid sequence variants, CADD is able to distinguish neutral from impactful variants across the entirety of the genome (Livesey and Marsh 2023). One of CADD’s unique features is that it is not restricted to SNPs in coding regions, gene-related information, or specific SNP categories, as it uses a large and diverse set of annotations. CADD was first applied to the human genome and subsequently CADD models were specifically designed for chicken, mouse and pig genomes (Groß et al. 2020a; Groß et al. 2018; Groß et al. 2020b; Kircher et al. 2014; Schubach et al. 2024). These different species-specific CADD models have shown that the CADD framework can be applied to different species provided that high-quality genome annotations are available (Boshove et al. 2023; Derks et al. 2021; Schubach et al. 2024; Speak et al. 2024; Wang et al. 2022).

While the CADD framework has been successfully applied to predict functional impact of variants in several species, it has not yet been extended to insects. The fruit fly *D. melanogaster* is a powerful genetic model, for which a wealth of genotyping data, high-quality genomes and single-nucleotide resolution annotations are available, including nucleotide conservation scores and regulatory domain annotations, which are necessary for the development of a species-specific CADD (Kim et al. 2024; Nunez et al. 2024; Öztürk-Çolak et al. 2024; Wangler et al. 2017). Moreover, the rich repertoire of gene editing approaches available for the *Drosophila* model system provides powerful opportunities to also functionally validate genotype-phenotype links at SNP-level, for example by applying CRISPR-Cas9 (Hoedjes et al. 2023; Perlmutter et al. 2024). The design of such functional genetic tests can be supported by functional impact predictions, which would make a species-specific CADD for the *D. melanogaster* a valuable tool to further our understanding of how genetic variation translates into phenotypic diversity by integrating diverse annotation with location-based information, which can provide crucial insights into the function(s) of SNPs.

Here, we introduce FlyCADD, the first adaptation of the CADD framework to an insect genome, applied to *D. melanogaster*. FlyCADD combines the available high-quality annotations into a robust and versatile tool for studying the functional impact of SNPs by providing impact prediction scores for all possible genomic variants in the genome of *D. melanogaster*. Combined annotations include conservation scores, gene structures, regulatory elements, GC content, and DNA secondary structure, among others. Combining in total 691 features into a single impact prediction metric, FlyCADD enables researchers to prioritize SNPs based on their predicted functional impact across different contexts and in any genomic region of the *D. melanogaster* genome.

FlyCADD offers a complementary approach when used alongside population data or experimental validation by refining the identification of potentially causal variants, thereby improving the understanding of genotype-phenotype relationships. Although FlyCADD, as a functional prediction tool for individual SNPs, cannot resolve the full complexity of genotype-phenotype relationships resulting from for example non-additive effect, epistasis or environmental effects, it can help researchers fine-scale their genotype-phenotype interpretations by distinguishing neutral from causal variants. Functional genomic studies to understand the impact of SNPs rather than genes in *D. melanogaster* are still rare, but the few precise genome editing studies done so far show unique and strong phenotypic effects of individual SNPs (Hoedjes et al. 2023; Mokashi et al. 2021; Perlmutter et al. 2024). SNPs identified in the genome of *D. melanogaster* natural populations, experimental strains or disease models can now be computationally studied based on predicted functional impact, with the potential to complement such experimental studies or prioritize SNPs for further (functional) studies. The FlyCADD model represents a significant step forward in integrating computational predictions into the study of genotype-phenotype link in *D. melanogaster*.

We describe the FlyCADD impact prediction model and demonstrate that impact predictions for SNPs can improve interpretation of candidate SNPs and experimental design. FlyCADD scores have been validated using experimentally tested point mutations with lethal outcome. To illustrate the usability of FlyCADD, we applied the FlyCADD impact prediction scores 1) to study naturally segregating SNPs, 2) to enhance interpretation of GWAS-identified SNPs, 3) to rank SNPs prior to functional studies, and 4) to improve experimental design for genome editing approaches. To facilitate variant impact analysis, we have made FlyCADD impact prediction scores available for all possible single nucleotide variants in the *D. melanogaster* Release 6 reference genome. Additionally, we provide scripts and pre-processed annotations for annotating novel variants. Both the precomputed scores and the locally executable pipeline can be accessed on Zenodo (https://doi.org/10.5281/zenodo.14887337).

## Methods

FlyCADD is a logistic regressor trained on 691 genomic features to assign an impact prediction score to any single nucleotide variant in the *D. melanogaster* genome. Impact refers to a mutation that on an evolutionary time scale is likely not tolerated at that position due to an associated fitness cost. FlyCADD is aimed at scoring the impact of single nucleotide variants; no other types of genetic variants were included in training, testing, or application of the model. An outline of FlyCADD development and application is illustrated in Figure 1. We build upon the existing CADD models for human, pig, and mouse by adapting the pipeline to fit *D. melanogaster* through incorporation of *D. melanogaster*-specific annotations, sequences, alignments, and the reference genome (Groß et al. 2018; Groß et al. 2020b; Kircher et al. 2014).

**Figure 1:**
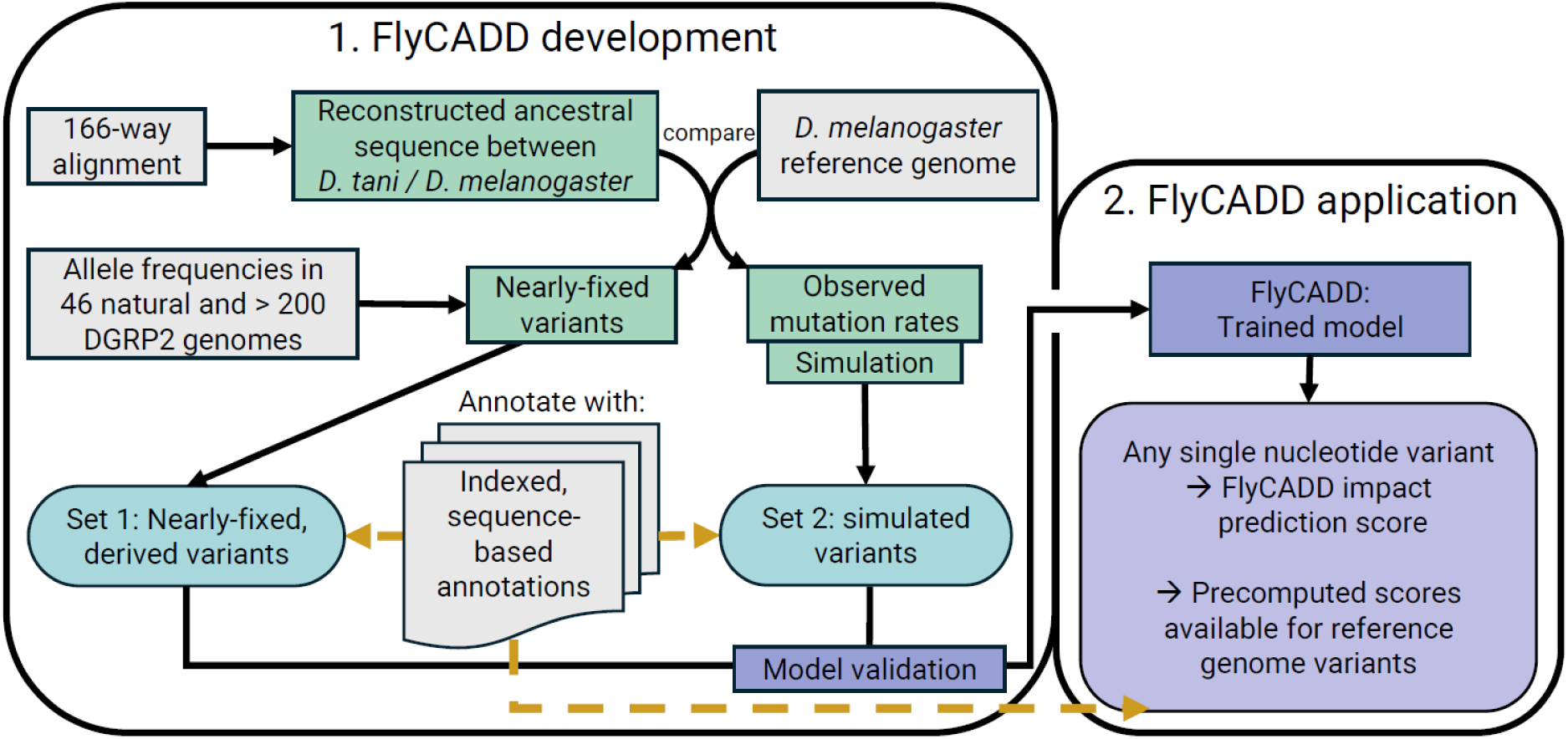
Overview of the FlyCADD pipeline consisting of two phases: FlyCADD 1) development and 2) application. Resources (grey) were used to extract information (green) that was applied to obtain the training datasets 1 and 2 (blue, rounded) for development of the FlyCADD model. These sets were annotated using 691 annotations (yellow dashed arrows). The model (purple) is trained on these this training dataset and can now be applied to any single nucleotide variant in the *D. melanogaster* genome, or precomputed scores can be used. To obtain impact prediction scores, variants of interest should be annotated (yellow dashed arrow) using the same annotations applied to the training datasets.

### Overview of FlyCADD model training

Briefly, the model consists of a logistic regressor trained to distinguish between two sets of training data: nearly-fixed derived variants - assumed to be depleted of impactful variants - and simulated variants - containing the full spectrum of impactful variants due to the absence of purifying selection. The FlyCADD model was created across chromosomes 2L, 2R, 3L, 3R, 4, and X based on the Release 6 reference genome (GCA_000001215.4). Ancestral sequence reconstruction was obtained from a Cactus whole-genome alignment of 166 *Drosophila* species in multiple alignment format (MAF)(Armstrong et al. 2020)(see File S1). This alignment was made available for FlyCADD development and represents an earlier version of the 298-way alignment described in (Kim et al. 2024). While both alignments were generated using the same methodology, the more recent version includes additional species and should be used for future analyses.

The set of derived variants was constructed by comparing the most recent common ancestor of *D. melanogaster* (melanogaster subgroup) and *D. tani* (montium subgroup) to the Release 6 *D. melanogaster* reference genome (GCA_000001215.4). Details of ancestral sequence reconstruction and extraction are described below (Methods “Obtaining training datasets”). Derived variants that are fixed or nearly-fixed (AF ≥ 0.9) in *D. melanogaster* populations across Europe and America with a different allele relative to the reconstructed ancestral genome are assumed to be depleted of deleterious variants due to purifying selection and belong to the derived variant set (proxy-benign variants). The second set of single nucleotide variants, containing simulated variants, was obtained by simulating *de novo* variants using mutation rates based on comparison between the ancestral and reference sequence. This set of simulated variants has not experienced natural selection and, therefore, compared to the derived set of variants, the simulated set is enriched for impactful alleles. In the following section (“Obtaining training datasets”), construction of these training datasets and variant simulation are described in more detail.

The second step in the CADD pipeline was to annotate the derived and simulated variants with a range of *Drosophila*-specific annotations and train the logistic regression model. The annotations included a total of 38 individual annotations such as sequence information, conservation scores, chromatin states, and combinations thereof, resulting in a set of 691 features (see File S2). The FlyCADD model was trained as a logistic regressor to classify annotated single nucleotide variants as belonging to either the derived or simulated class. The logistic regressor returns the probability of a variant belonging to the set of simulated variants and a probability of a variant belonging to the set of derived variants, which sum up to 1. For each single nucleotide variant on the reference genome, the impact prediction score representing the probability of the variant belonging to the simulated variants class is readily available. The impact score is a measure for predicted functional impact to fitness between 0 (low functional impact) and 1 (high functional impact). All annotations and impact prediction scores are 1-based. The pipeline for FlyCADD development can be found on GitHub (https://github.com/JuliaBeets/FlyCADD).

### Obtaining training datasets

A Cactus 166-way alignment of *Drosophila* species was provided for ancestral sequence reconstruction, prior to publication of the 298-way alignment (Kim et al. 2024). The Cactus alignment software reconstructs ancestral genomes for each node, calling individual bases as the maximum likelihood estimate from a Jukes-Cantor substitution model (Armstrong et al. 2020). The 166-way alignment including ancestral sequence reconstructions used in this work is available on Zenodo (https://doi.org/10.5281/zenodo.14887337). The ancestral sequence was obtained by extracting the reconstructed sequence on the sense strand relative to coordinates in Release 6 of the reference genome for the most recent common ancestor between *D. melanogaster* and *D. tani* within the 166-way alignment (see File S1). This ancestral node was chosen as the reconstructed sequence was the largest, with 16.4 % of the reference sequence nucleotides having a corresponding nucleotide in the ancestral sequence, whereas the rest of the ancestral sequence consists of gaps. Additionally, the chosen ancestral sequence showed no excessive bias towards specific chromosomes or coding sequences. Considering these aspects of bias is important to ensure that derived and simulated variants are as evenly distributed across the genome as possible. A more detailed description of ancestral sequence reconstruction can be found in File S1.

To construct the set of derived variants, alleles in the reference genome differing from that of the ancestral genome were identified. Subsequently, to ensure that the set of derived variants included (nearly) fixed variants, variants with an allele frequency below 0.9 were removed. The *D. melanogaster* populations used to define allele frequencies included 46 genomes from 12 *D. melanogaster* populations in Europe and America (Rech et al. 2022) and > 200 Drosophila Genetic Reference Panel 2 (DGRP) lines (Mackay et al. 2012b). This resulted in a set of 2.733.695 derived variants, which is assumed to contain mainly benign or neutral variants.

An equal number of variants was obtained through simulation using the variant simulator of the original CADD model to create a set of simulated variants assumed to be enriched for impactful variants compared to the set of derived variants (Kircher et al. 2014). Mutation rates and CpG sites were calculated per chromosome based on comparison between the reconstructed ancestral sequence and the reference genome and were applied to simulate variants that are absent in the reference genome but could have naturally occurred based on mutation rates in the absence of purifying selection and free from the influence of biological selection. The final mutation rates applied in the simulation can be found on GitHub (https://github.com/JuliaBeets/FlyCADD). The set of simulated *de novo* mutations was then randomly trimmed to match the number of derived variants.

The final numbers of derived and simulated variants per chromosome are summarized in Table 1. The Y chromosome was excluded from model training due to the lack of a sufficiently reconstructed ancestral sequence required to construct the datasets of derived and simulated variants. In total, the training dataset contained 5.467.390 single nucleotide variants, equally divided over the set of derived variants and the set of simulated variants.

**Table 1:** Final numbers of derived and simulated SNPs used in training and testing the logistic regression model.

|  | Derived | Simulated |
| --- | --- | --- |
| **Chromosome 2L** | 594.922 | 662.282 |
| **Chromosome 2R** | 471.645 | 435.022 |
| **Chromosome 3L** | 563.599 | 577.720 |
| **Chromosome 3R** | 658.177 | 659.379 |
| **Chromosome 4** | 25.216 | 15.959 |
| **Chromosome X** | 420.148 | 383.345 |
| **Total** | 2.733.695 | 2.733.695 |

### Annotations

The training dataset was annotated with a total of 691 features (38 distinct annotations and relevant combinations thereof). The Ensembl Variant Effect Predictor (VEP, v105.0) provided 13 sequence-based annotations in offline mode (McLaren et al. 2016). The proportion of variants per VEP consequence in the derived and simulated variant datasets can be found in File S3. These were supplemented with Grantham-scores, secondary structure predictions, repeat annotations, regulatory elements, PhyloP, PhastCons, GERP, chromatin state, miRNA, coding regions, transcription factor binding sites, and *cis*-regulatory motifs. Details for all features can be found in File S2, and the sources of these annotations are described below.

Grantham scores were incorporated based on the Grantham-matrix that annotates the chemical difference induced by the amino acid change upon variation (Grantham 1974). Secondary structure predictions for the reference genome were obtained using the *getshape*() function from DNAshapeR (Chiu et al. 2016). The included features were minor groove width (MGW), roll, helix twist (HelT), propeller twist (ProT), and electrostatic potential (EP).

Coordinates for eight repeat types (DNA repeat elements, low complexity repeats (LCR), long interspersed nuclear elements (LINE), long terminal repeat elements (LTR), rolling circle (RC), satellite repeats, simple repeats, and RNA repeats) were retrieved from the RepeatMasker Track of the UCSC Genome Browser (v460, last updated 2014-08-28)(Perez et al. 2025).

ReMap records of regulatory elements, PhyloP base-wise and PhastCons element conservation scores, were obtained from the UCSC Genome Browser (v460, last updated 2014-08-28)(Perez et al. 2025). ReMap records were incorporated in FlyCADD as the density of records per position. PhyloP and PhastCons scores were based on different multi-species alignment sizes: Multiz alignments of 27 insects (PhyloP/PhastCons) and 124 insects (PhyloP124/PhastCons124). Additionally, PhyloP scores (PhyloP_all) and GERP-like scores (GerpRS_all and GERPN_all) were computed from the 166-way Cactus alignment with PHAST (Hubisz et al. 2010), using the implementation of the tools in the halPhyloPMP.py script provided as part of the HAL toolkit (Hickey et al. 2013), excluding the *D. melanogaster* sequence to avoid bias.

Chromatin states for BG3 and S2 cells, miRNA encoding sequence coordinates, and coordinates for coding sequences were retrieved from FlyBase (Release FB2023_03)(Öztürk-Çolak et al. 2024). Coordinates for *cis*-regulatory motifs, predicted *cis*-regulatory motifs, and transcription factor binding sites were retrieved from REDfly (v9.6.2)(Keränen et al. 2022).

Several annotations were combined with VEP annotations to create the final set of 691 features (see Table 2 in File S2). These include all combinations of amino acid changes, variant consequences, conservation scores, ReMap scores, and distances to coding sequences. All features were scaled by their standard deviation based on variation within the training dataset. In the final annotated dataset, derived variants were annotated with “1” and simulated variants with “0”.

### Logistic regression model

To select the optimal L2 regularization parameter for the logistic regression model, we used repeated random sub-sampling validation. The data was split five times into 90% training and 10% testing subsets for each potential value of L2 (0.01, 0.1, 1.0, 10.0, 100.0). For each split, a model was trained and evaluated. The mean accuracy across splits was used to select the best L2 penalty. The final Turi Create Logistic Classifier (v6.4.1) was trained on 90% of the annotated dataset, containing variants in two categories—“derived” and “simulated”-, reserving the remaining 10% for model testing, with L2 penalization set to 1.0, using the Newton solver, and a maximum of 100 iterations with *tc.logistic_classifier.create*(). A trained model was obtained for the full genome, excluding the Y chromosome due to its absence in the training dataset. The model assigns weight to each feature after training, representing the predictive power of the feature. The directionality of the contribution cannot be interpreted with biological meaning regarding functional impact and rather stems from the encoding of the features themselves. The model accuracy was evaluated using a built-in testing functionality of the logistic regression function applied to the held-out 10 % of the training dataset, ensuring that none of these SNPs were used for training.

### Computing FlyCADD scores

The FlyCADD model can be applied to any given variant on the *D. melanogaster* genome that is annotated with the same set of features and scaled by the standard deviation per feature within the training data. With *loaded_model.predict*(), FlyCADD generates a raw probability score for each variant, indicating its likelihood of belonging to the simulated class. FlyCADD scores range from 0, indicating no predicted functional impact, to 1, indicating high predicted functional impact. The FlyCADD model was applied to all possible single nucleotide variants on the *D. melanogaster* reference genome autosomes and X chromosome, resulting in posterior probabilities for each variant belonging to the simulated variants set and thus being impactful. Since these precomputed impact prediction scores for all possible genomic variants include those genomic variants not observed in natural populations, predictions are unbiased by allele frequencies, function or location of known variants and solely based on variant annotations and model training. The precomputed impact prediction scores, trained model and scripts to score novel variants are readily available on Zenodo (https://doi.org/10.5281/zenodo.14887337).

### Variant impact at codon positions

Nucleotide positions within codons differ in biological importance, providing an opportunity to evaluate the impact prediction scores (Crick 1966). Genome-wide codon positions were retrieved from Ensembl (v111) and filtered for unique transcripts on the sense strand, resulting in transcripts of 6.894 genes. Genes with multiple transcripts were excluded to avoid interaction effects. The distributions of minimum FlyCADD scores within a codon were compared per position amongst all transcripts and within each gene using the Mann-Whitney U-test. Statistical analyses were performed using scipy.stats (v1.12.0) in Python3, with Bonferroni corrected p-values (Virtanen et al. 2020).

### Impact predictions of lethal, chemically induced point mutations

To assess the performance of FlyCADD on empirically tested point mutations, we retrieved data on point mutations that were chemically induced using ethyl methanesulfonate (EMS) with lethal phenotypic outcome using QueryBuilder (FlyBase FB2025_02). This resulted in a total of 2202 records of which 2168 variants had a known position and base change. After excluding duplicates, FlyCADD scores were extracted and analyzed for 2118 unique point mutations.

### Allele frequency of natural variants

To investigate the distribution of FlyCADD scores for naturally occurring SNPs with differing allele frequency, positions and allele frequencies for over 4.8 million SNPs were obtained from the VCF file (version dest.all.PoolSNP.001.50.24Aug2024) of the *Drosophila* Evolution over Space and Time (DEST) 2.0 resource, the most comprehensive genomic resource for *D. melanogaster* genomes worldwide (Nunez et al. 2024). Only samples with quality filtering label “PASS” (n = 529) were included in this analysis to avoid inclusion of low-quality variants.

## Results

FlyCADD predicts the functional impact of SNPs in *D. melanogaster*. Impact scores were precomputed for all possible single nucleotide variants on the Release 6 reference genome (excluding the Y chromosome), with higher scores reflecting greater predicted functional impact of the SNP. FlyCADD scores range from 0 to 1, where lower scores suggest a benign or neutral effect, while higher scores (e.g., > 0.60) indicate potential functional impact of the SNP. These scores allow researchers to rank SNPs based on their potential impact and assess the functional relevance of candidate SNPs. Here, we evaluate these scores based on the accuracy metrics of the model, contributions of different annotations, biologically relevant impact predictions, point mutations with known phenotypic effect and we show the tool’s applicability across diverse use cases.

### High accuracy achieved by training the logistic regression model

The performance of the FlyCADD logistic regression model was evaluated using a held-out 10 % of the training dataset (see Methods). The overall performance for FlyCADD is ∼ 0.83 (area under the ROC curve) with an accuracy of ∼ 0.76, reflecting the proportion of correct predictions. It is important to note that FlyCADD’s accuracy evaluation relies on a binary classification approach, where variants in the testing dataset were labeled strictly as 0 (neutral) or 1 (impactful), similar to the variants in the training dataset. However, functional impact is a continuous spectrum, and some variants labeled as 0 may still have (minor) effects. A prediction could be interpreted as misclassification during testing despite reflecting a biologically meaningful ranking, therefore, the accuracy of the model is likely to be higher when considering an impact gradient or using validation datasets for model evaluation.

### Key predictors are combined features

The strength of the CADD framework lies in its ability to integrate both individual and combined annotations, allowing for more precise distinction between variants. FlyCADD incorporated 38 individual annotations and combinations thereof (see File S2). Out of the total 691 features used for annotation by FlyCADD, 581 were combinations of individual annotations. The model assigns weights to these features, reflecting their contribution to the impact prediction. The weight indicates the predictive power of a feature within the logistic regression model. Ultimately, the impact prediction score is a combined score, considering all 691 features of a variant. The weights for all these features can be found in File S3.

Figure 2 shows the twenty features with the highest predictive power within the model, i.e. which features most strongly influence the model’s ability to predict whether a variant is neutral or benign, or impactful. These features enhance predictive power but do not independently determine the predicted functional effect of a SNP. When considering the twenty most influential features in the model (Figure 2), it is visible from the composed feature names that most key features are combinations of annotations. These combinations involve the annotations on conservation scores (“_PhyloP”), proximity to coding sequence (“_relCDSpos”) and consequences (synonymous and non-synonymous, “SN_”/”NS_”). The individual annotations amongst the most influential features of FlyCADD are position within the protein (“protPos”), consequences stop loss, non-synonymous, synonymous and stop gain (“Consequence_”), stop codon as alternative allele (nAA_*) and conservation score (PhyloP).

**Figure 2:**
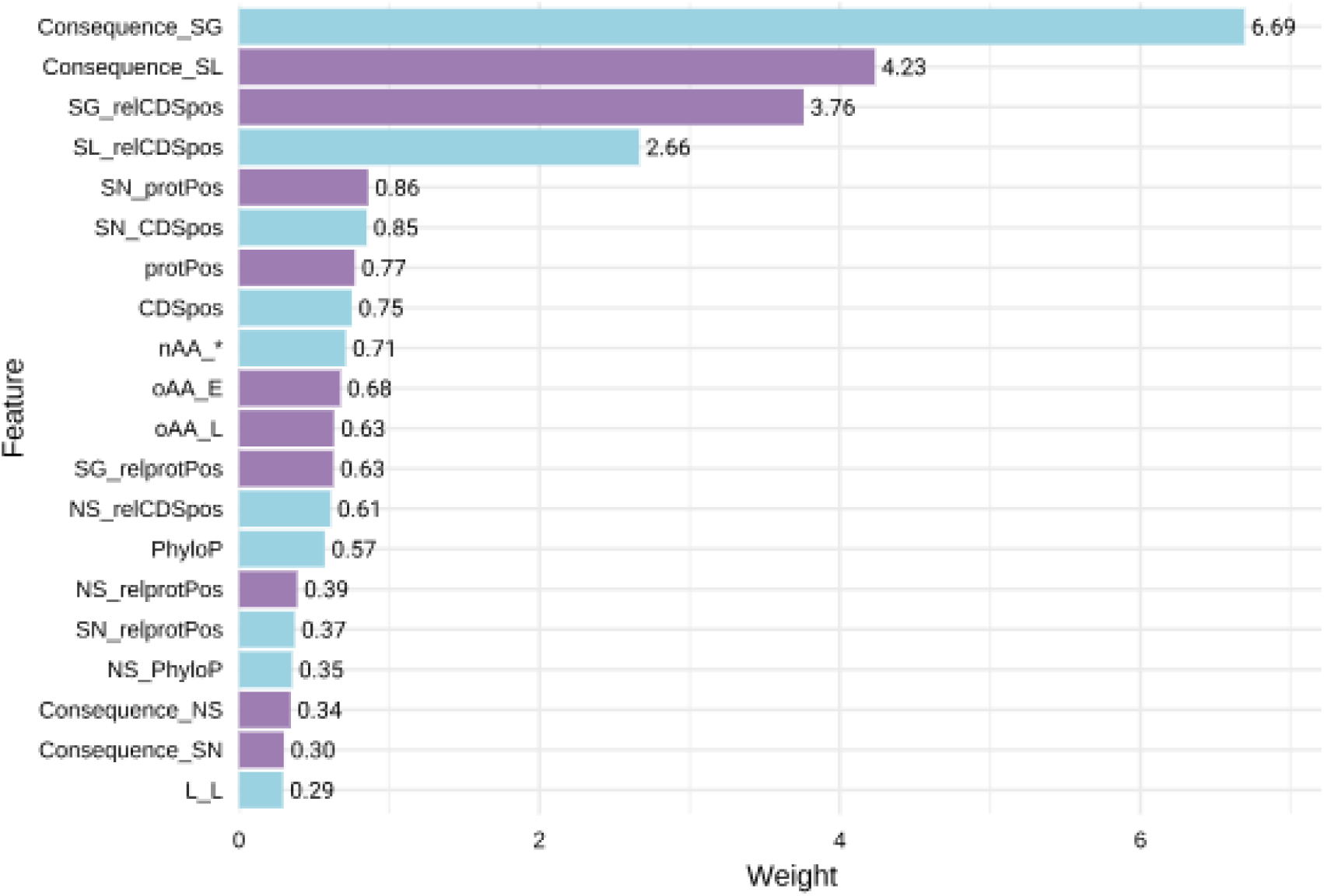
Feature weights for the twenty features with most predictive power. The color indicates a negative (blue) or positive (purple) weight as computed by the logistic regression model. Features names “Consequence_” reflect the predicted VEP consequence derived from the Ensembl Variant Effect Predictor. Abbreviations: (rel)CDSpos: (relative) position within the coding sequence; (rel)Protpos: (relative) position within the protein; E: glutamine; L: leucine; nAA: alternative amino acid; NS: non-synonymous substitution; oAA: original amino acid; SG: stop-gain; SL: stop-loss; SN: synonymous; *: stop codon. Detailed explanations of all features can be found in File S2.

Moreover, the combined features in the model capture complex interactions that enhance predictive performance beyond what individual annotations can provide. For instance, features related to stop codon loss and gain, when considered together with their position within the coding sequence, rank among the top predictors of variant impact. This reflects the biological importance of these polymorphisms, as changes affecting protein length and their location within the protein sequence strongly influence functional consequences (Lee and Reinhardt 2012). Overall, these twenty features demonstrate the highest predictive power in distinguishing impactful variants, underscoring the effect of integrating multiple annotations into a functional impact score.

### Genome-wide FlyCADD scores reflect biological relevance of polymorphisms

Next, we analyzed the precomputed FlyCADD scores for all possible single nucleotide variants on the *D. melanogaster* reference genome. Importantly, this set does not represent naturally occurring variants but rather includes all theoretically possible variants relative to the reference genome. The genome-wide distribution of FlyCADD scores shows that predicted high-impact variants (> 0.95) are the most prevalent, while predicted low-impact variants (0.00 – 0.1) are the least frequent, a pattern that is consistent across chromosomes (Figure 3). The limited number of low-impact scores (< 0.1) likely reflects that many of the theoretically possible SNPs in this dataset do not actually segregate in natural populations. For example, the DEST genomic resource has identified ∼ 5 million SNPs segregating in natural populations, whereas this set which includes all possible variants on the *D. melanogaster* reference genome contains > 400 million variants, most of which are not occurring naturally. These non-natural SNPs are likely to have higher functional impact and thus receive higher FlyCADD scores, resulting in a low proportion of variants with low impact scores in this dataset. Variants with such low predicted impact reflect a pattern of annotations by FlyCADD similar to the nearly-fixed, derived variants, as these are the proxy-benign variants in training. The theoretically possible SNPs in the set of all possible SNPs on the reference genome do not reflect such patterns as they are likely not variants that will become nearly-fixed or occur at all, and this results in high predicted impact for most variants in the dataset.

**Figure 3:**
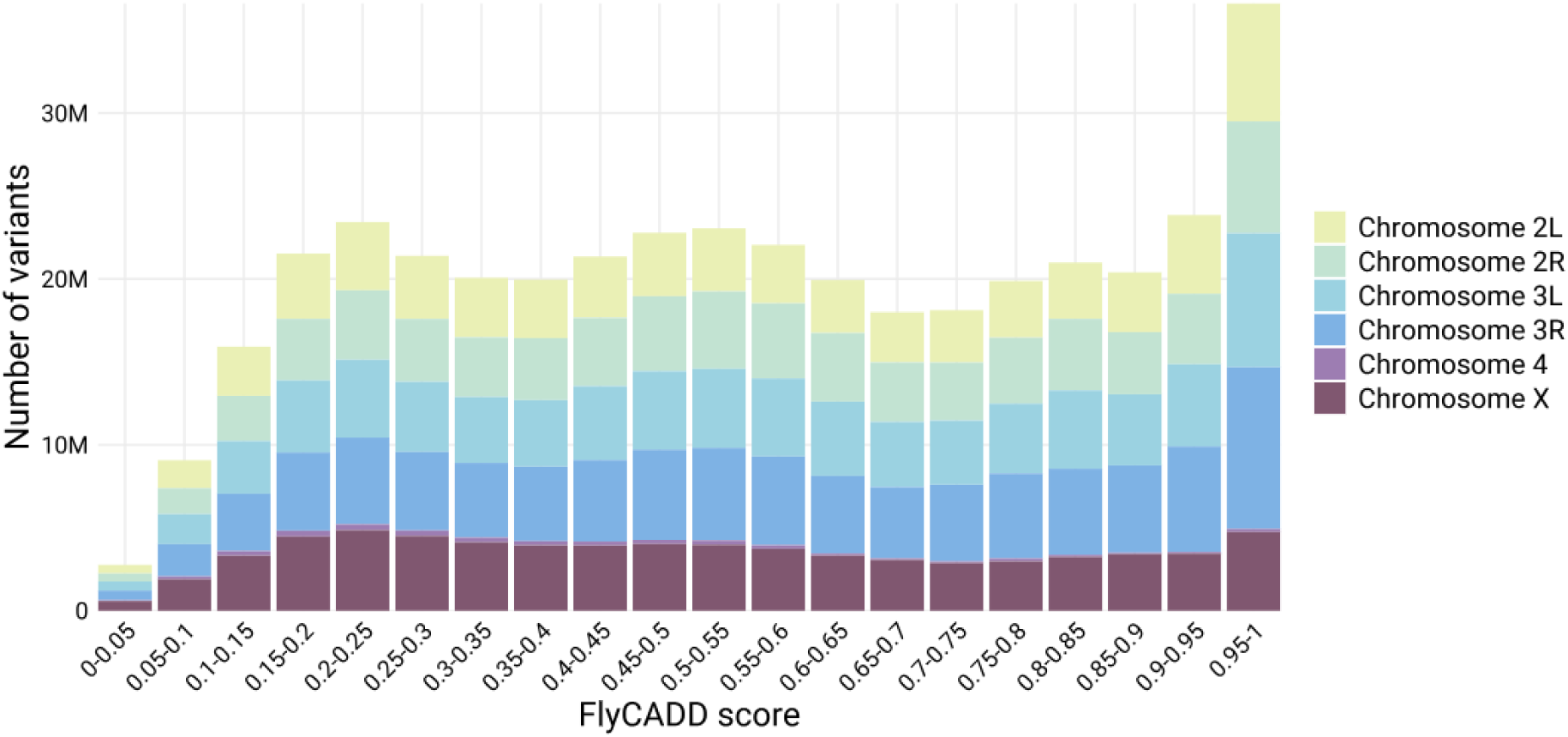
Distribution of FlyCADD scores in the set of precomputed scores for all possible single nucleotide variants on chromosomes 2L, 2R, 3L, 3R, 4 and X of the *D. melanogaster* reference genome.

Subsequently, we assessed whether FlyCADD scores vary among variants when categorized by predicted consequence (based on VEP), such as gain of a stop codon, intronic or synonymous variants, to provide insights into FlyCADD’s ability to predict functional impact across classes of variants (Figure 4). For example, it is known that non-synonymous mutations and stop codon polymorphisms are impactful as they alter the protein. In line with our hypothesis, synonymous (SN) variants have lower FlyCADD scores, whereas non-synonymous (NS) variants have higher scores, potentially due to their amino acid changing properties (Figure 4).

**Figure 4:**
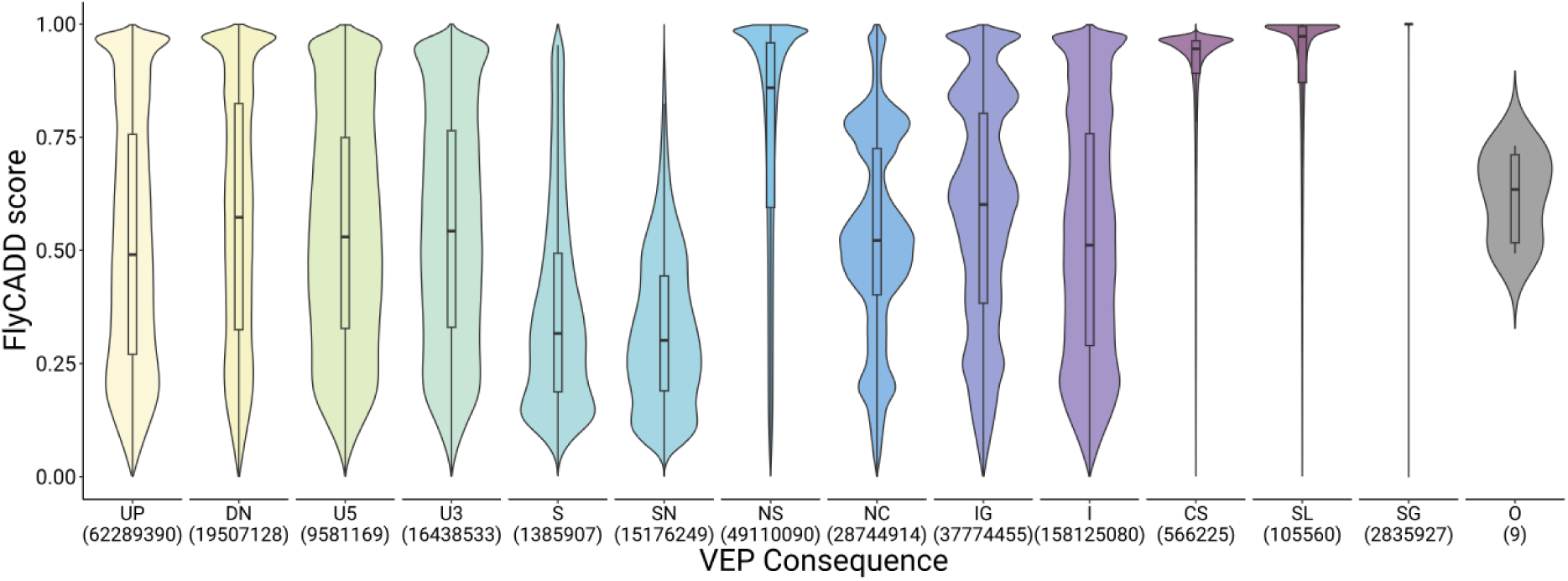
Distribution of FlyCADD scores for all possible single nucleotide variants on the *D. melanogaster* reference genome categorized by predicted consequences from VEP. Violin plots show score density per consequence; the median and interquartile range are indicated. UP: upstream; DN: downstream; U5: 5 prime UTR; U3: 3 prime UTR; S: splice site; SN: synonymous; NS: non-synonymous; NC: non-coding exon change; IG: intergenic; I: intronic; CS: canonical splice; SL: stop loss; SG: stop gain; O: other.

However, there are SN variants present in the dataset with higher predicted impact and, alternatively, NS variants with lower predicted impact are present (Figure 4). The range of predicted impact for both these types of variants is large and overlapping. Therefore, applying FlyCADD to NS or SN point mutations can help refine interpretation of the functional impact of the variants within these consequence classes instead of solely relying on the individual VEP consequence prediction. Variants with other consequences showed a broader FlyCADD score distribution compared to SN or NS variants, reflecting the additional information captured by integrating 691 features. For example, intergenic and non-coding variants exhibited widely spread scores, highlighting how combining diverse annotations into a single metric enhances interpretation, especially in less characterized genomic regions where single annotations are often less informative. FlyCADD thus enables meaningful ranking of SNPs across the genome, including those in regions with limited prior functional insight.

### FlyCADD scores refine impact prediction within gene structure

Genes are composed of distinct functional regions, such as introns, exons and transcription factor binding sites. We found that FlyCADD scores reflect the underlying genomic arrangement by recognizing, amongst others, exonic variants with generally higher FlyCADD scores compared to intronic variants when considering all possible single nucleotide variants within genes. This is illustrated in Figure 5 for the gene *white* where, as expected, variants in exons generally receive higher FlyCADD scores compared to intronic variants. However, both regions contain SNPs that cover the full range of FlyCADD scores. Additionally, naturally occurring variants in such regions exhibit a wide spread of predicted impact (Figure 5). By evaluating SNP impact using combined annotations rather than relying solely on gene structure, FlyCADD enables functional impact prediction at single-nucleotide resolution rather than considering all variants within a functional domain as equivalent. Traditional approaches often infer SNP impact based on proximity to genes or annotated domains, overlooking functional differences between variants within the same region. FlyCADD allows researchers to distinguish between neutral and functionally relevant variants within these functional regions.

**Figure 5:**
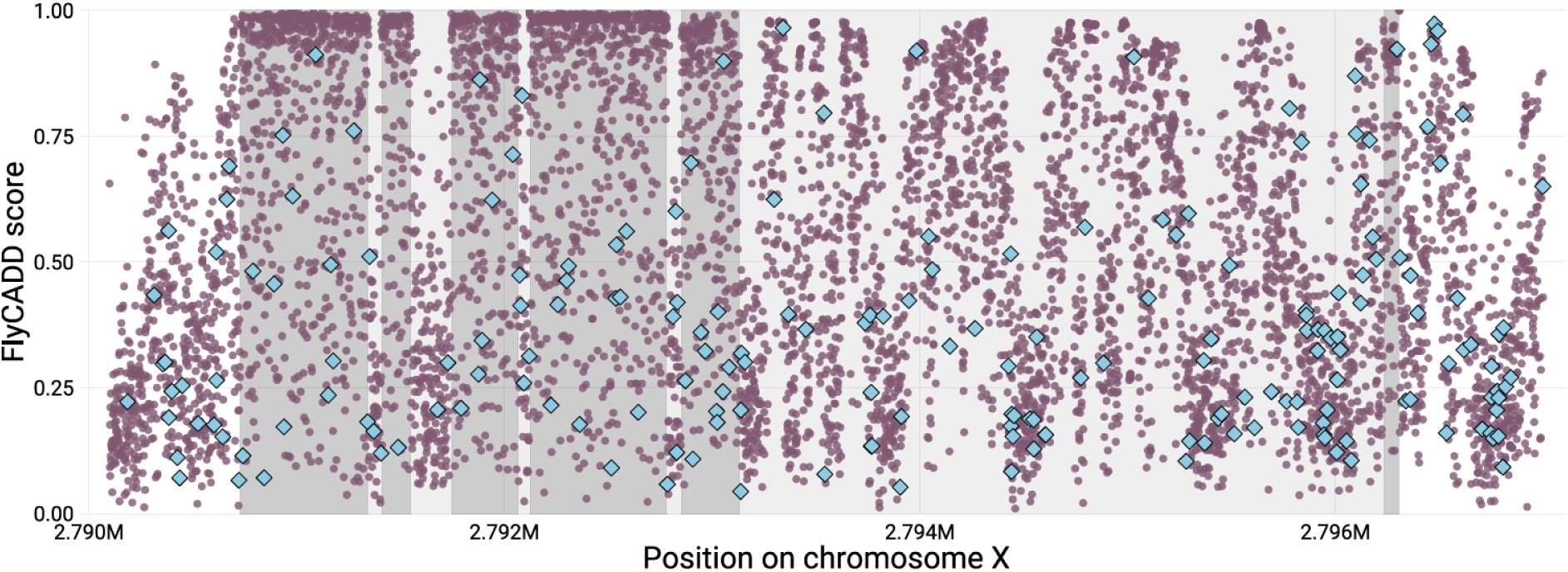
Mean FlyCADD scores (purple) per position for all possible single nucleotide variants in the gene *white* on chromosome X, including 500 bp upstream and downstream of the gene. Exons of the gene are marked dark grey, introns marked in light grey. Naturally occurring variants retrieved from DEST2.0 are indicated in blue.

### Nucleotides at each codon position show differing predicted impact

To investigate the performance of FlyCADD impact predictions at single nucleotide resolution, we examined variation in FlyCADD scores across codon positions, which differ in functional importance due to their unique roles in DNA secondary structure, transcription, translation and mutation accumulation (Crick 1966). The third codon position is generally least impactful, as variation at this site rarely results in amino acid changes (Crick 1966).

We compared the distribution of minimum FlyCADD scores of all possible variants on the three codon positions across 6.894 genes (Figure 6a), using the lowest-scoring variant at each position to provide a conservative estimate of functional impact and avoid bias from outliers. Variants at the first and second codon position both have significantly higher FlyCADD scores compared to the third nucleotide. This is in line with expectations since the third codon position is often referred to as the “wobble” nucleotide, where base pairing is less strict and amino acids are less likely to be altered by changes in the third nucleotide (Crick 1966). The third nucleotide has the least functional impact, followed by the first nucleotide, whereas the second nucleotide has the most functional impact. Additionally, we compared the distribution of FlyCADD scores of the three codon positions per gene (Figure 6b).

**Figure 6:**
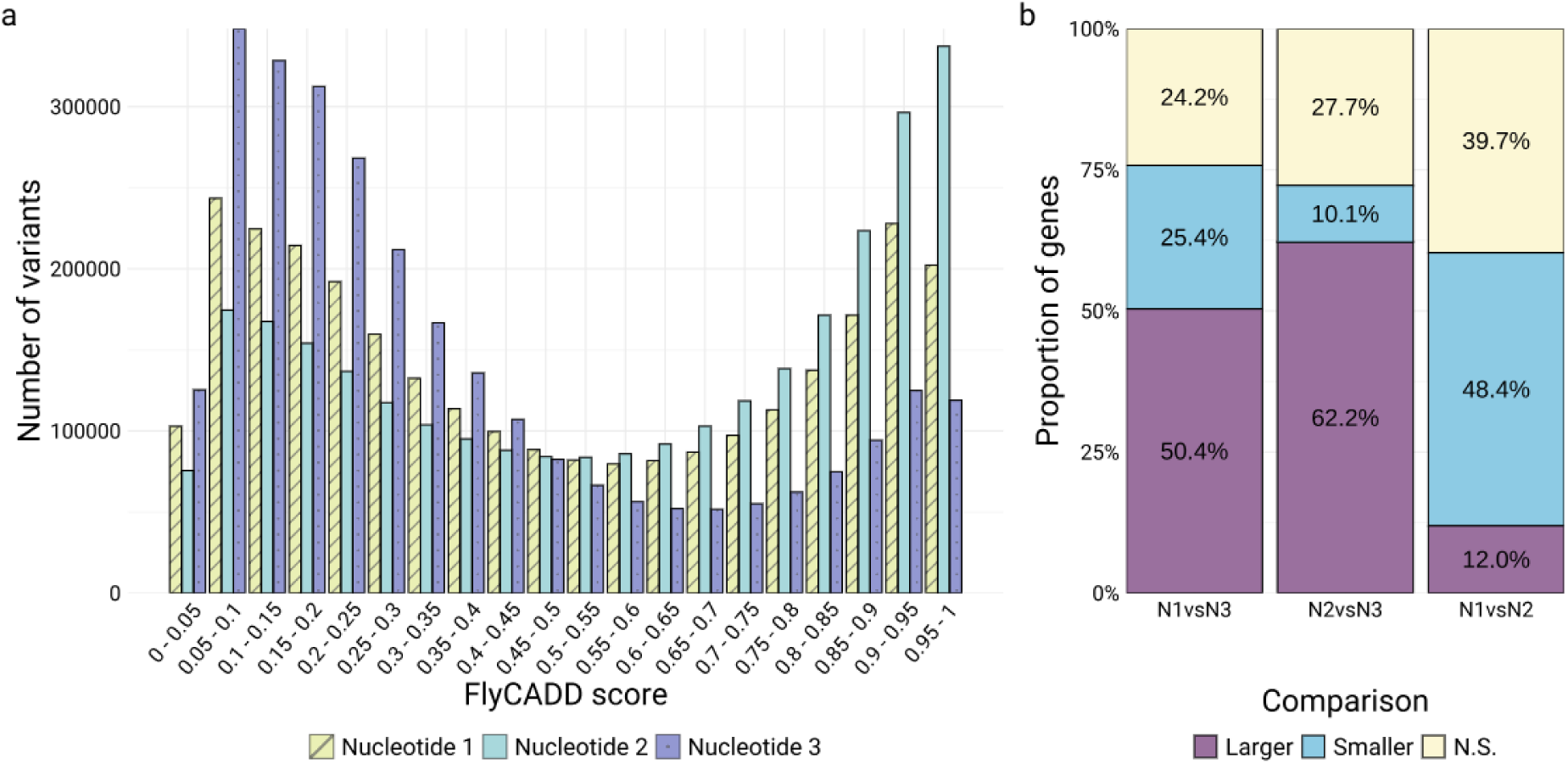
Distribution of minimum FlyCADD scores of variants at each of the positions within the codon;(a) distribution across 6.894 transcripts and (b) FlyCADD scores at each nucleotide in a codon compared within each gene. The proportion of genes where the distribution of FlyCADD scores significantly differed between two nucleotide positions after Bonferroni corrected Mann-Whitney U tests is shown in panel b.

Analysis shows that the first nucleotide in each codon has significantly higher FlyCADD scores compared to the third nucleotide in 50.4 % of the genes, whereas the second nucleotide scores higher compared to the third nucleotide in 62.2 % of the genes (Figure 6b). Therefore, the third nucleotide has the lowest FlyCADD scores in the majority of genes. The first codon position was scored significantly higher by FlyCADD compared to the second nucleotide in only 12 % of the genes, indicating that the second codon position is more important than the first in many genes (48.4 %). Taken together, the impact predictions indicate that nucleotides at the third codon position are least functionally important, followed by those at the first, with nucleotides at the second position having the greatest impact. This is consistent with observations across all genes (Figure 6a) and indicates that FlyCADD scores reflect biologically expected patterns of functional impact at the single-nucleotide level.

### Lethal point mutations predicted as high impact variants

FlyCADD was developed without the use of experimental phenotype annotations, nor was it trained on known impactful mutations. To evaluate the performance of FlyCADD to predict the impact of mutations with significant phenotypic impact, we applied FlyCADD to EMS-induced point mutations with known lethal outcome retrieved from FlyBase. A total of 2118 unique point mutations with lethal outcome were identified and FlyCADD identified 93.3 % of these mutations as impactful with impact prediction scores above 0.6 (Figure 7). 88.6 % even had a predicted impact of 0.8 and higher, strongly indicating functional impact which corresponds with their involvement in lethality. These variants with high phenotypic severity are recognized as impactful mutations by FlyCADD indicating that the evolutionarily informed framework has a high accuracy to assess the impact of functionally tested point mutations. Several variants with a low predicted impact are also present in this dataset and might indicate variants lethal under distinct circumstances such as in specific environmental or genetic backgrounds. For example, two point mutations in the gene *kon* receive a FlyCADD score of 0 indicating no functional impact. However, they are associated to the lethal phenotypic outcome in a specific genetic background; a T to A point mutation at chr2L:18492868 is lethal when occurring with a four base pair deletion and the point mutation at chr2L: 18494189 is lethal when co-occurring with other point mutations in this gene (Estrada et al. 2007; Schnorrer et al. 2007). FlyCADD scores indicate which of these point mutations is likely to be the causal factor.

**Figure 7:**
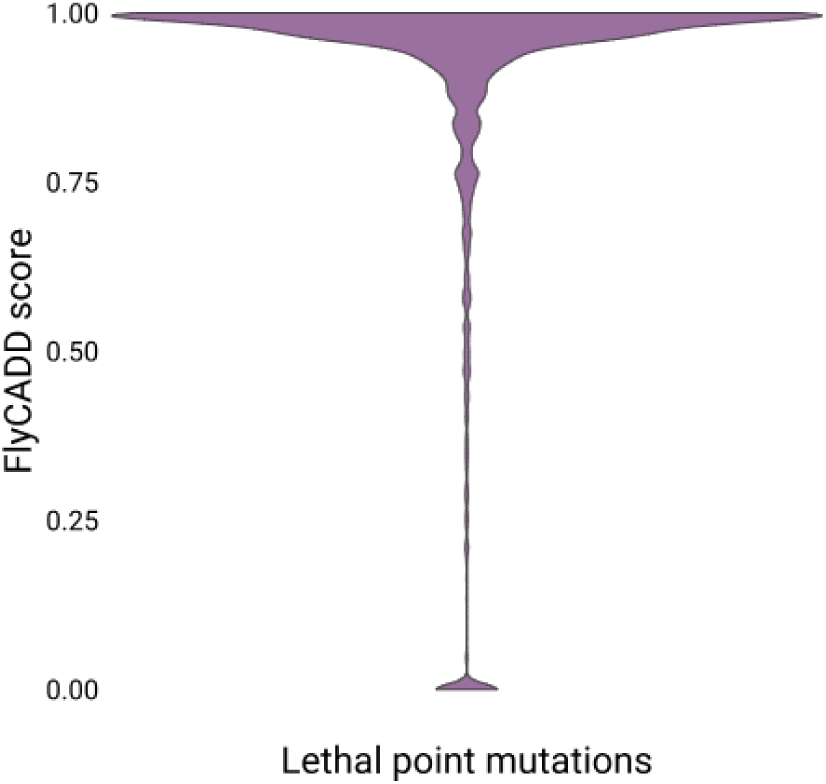
Distribution of FlyCADD scores for 2118 chemically induced point mutations with lethal outcome retrieved from FlyBase.

### Use cases for FlyCADD scores

Approaches such as GWAS and E&R often yield large numbers of candidate SNPs associated with a phenotype of interest, but these often include false positive and false negative hits due to linkage or technical limitations, such as statistical power (Wangler et al. 2017). In addition, experimental validation of candidate SNPs is costly, time-consuming and not yet feasible for large panels of SNPs. The FlyCADD model generates impact prediction scores reflecting a fitness-reducing effect of SNPs, making it a promising tool for genetic research by facilitating the prioritization and interpretation of identified SNPs across a range of studies and topics. Here, we outline four key applications of FlyCADD to study *D. melanogaster* genomic variation at single nucleotide resolution, each illustrated with a specific example demonstrating the utility of impact prediction scores in various research contexts.

#### Impact of SNPs in natural populations of D. melanogaster

Natural populations of *D. melanogaster* exhibit extensive genetic diversity, with large numbers of segregating alleles continuously being identified (Nunez et al. 2024). While this natural variation is a driver of adaptation, genetic variation without functionally relevant impact is likely to circulate in populations as well. Methods for identification of functional loci from population genomic resources are often not focused on the functional impact of individual variants, but rather on sets of SNPs or nearby genomic regions (Harr et al. 2002; Jha et al. 2015). The FlyCADD model can filter out hitchhiking loci or variants with negligible functional effects from population genomics studies, and identify naturally occurring, high-impact SNPs. Furthermore, FlyCADD scores offer a basis for identifying ranges to classify variants by their potential impact, providing insights into the tolerability of variation within natural populations.

The genomic resource “*Drosophila* Evolution over Space and Time” (DEST) 2.0 contains high-quality whole-genome sequence information of 529 *D. melanogaster* populations worldwide (Nunez et al. 2024). When applying FlyCADD to the ∼4.8 million naturally segregating SNPs from this dataset, we observed a mean FlyCADD score of 0.45, with an increased mean impact prediction score (0.50) observed for rare alleles (AF < 0.05)(Figure 8). When excluding rare alleles, the mean FlyCADD score is 0.36. This data is indicative of a higher predicted functional impact for rare alleles. Naturally occurring variants are showing a distribution of FlyCADD scores across the full range (0 – 1), irrespective of AF. Outliers with FlyCADD scores exceeding 0.80 (high impact) are more commonly observed among rare alleles, while becoming less frequent as AF increases. The presence of both low and high impact variants highlights the potential value of FlyCADD in differentiating functionally relevant variants from neutral loci within population genomic resources like DEST.

**Figure 8:**
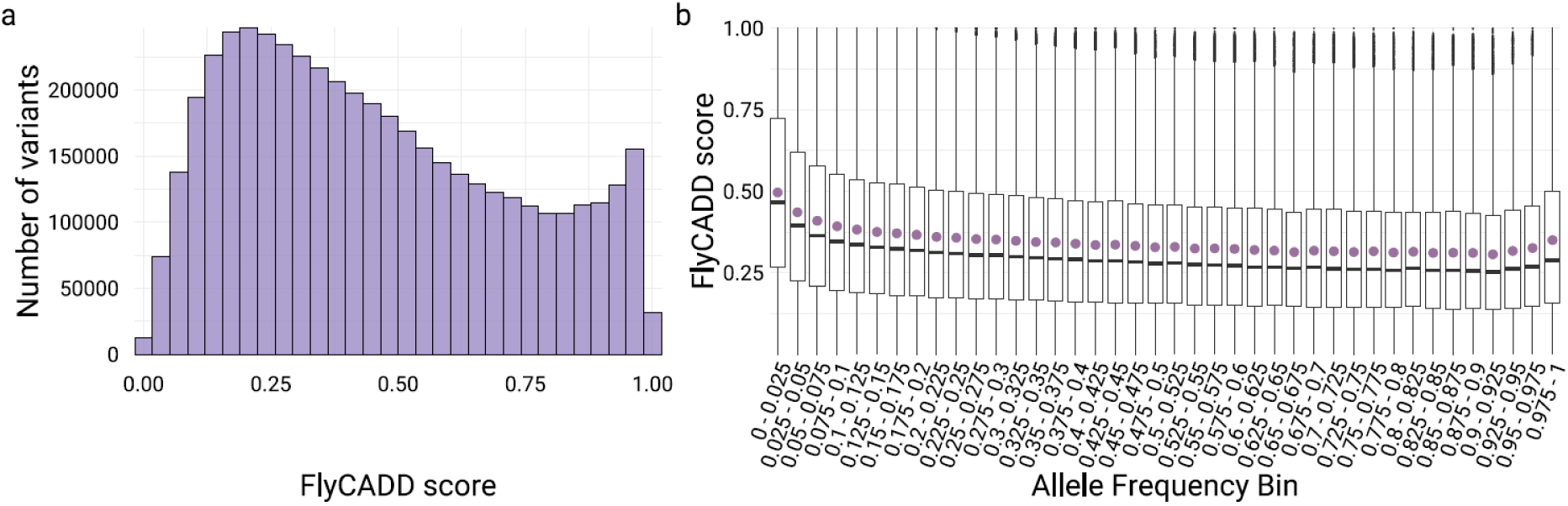
FlyCADD scores of naturally occurring SNPs, identified in the DEST2.0 genomic resource. (a) Distribution of FlyCADD scores and (b) boxplots showing the distribution of FlyCADD scores across allele frequency bins (bin size 0.025). Allele frequency represents the frequency of the non-reference allele (as mapped to the reference genome). The boxplots display the median, interquartile range and spread of FlyCADD scores. The mean FlyCADD score for variants per allele frequency bin is marked purple.

Based on the distribution of FlyCADD scores of naturally occurring SNPs (Figure 8), we propose a range for interpreting functional impact of naturally occurring SNPs. Variants with scores between 0 and approximately 0.40 appear to be generally tolerated in natural populations as they are frequently present in natural populations, whereas those with scores above 0.60 emerge as candidates for further study due to their high potential for functional impact. However, the degree to which functional impact influences populations may be shaped by ecological and selective pressures.

#### FlyCADD-based ranking compared to p-value ranking of SNPs from GWAS

Currently, p-values are the most common metric used for prioritization of phenotype-associated candidate SNPs resulting from GWAS. Solely relying on p-values for ranking GWAS SNPs is prone to confounding factors such as linked variants, statistical limitations and environmental factors, which are well-known to influence these p-values (Chen et al. 2021; Fadista et al. 2016; Korte and Farlow 2013). Therefore, p-values give a first insight into candidate loci, but identification of causal SNPs from these results is troublesome and subject to varying cut-off values (Cano-Gamez and Trynka 2020; Korte and Farlow 2013). As we demonstrate below, FlyCADD scores can complement the p-value ranking derived from association studies by providing an impact prediction score per SNP and potentially offer additional insight into evolutionary fitness consequences of candidate SNPs.

A GWAS study by Katzenberger et al. provides an opportunity to apply FlyCADD scores in addition to p-values for assessment of functional impact of SNPs (Katzenberger et al. 2015). The study investigated the genetic basis of mortality after induced traumatic brain injury in DGRP lines, identifying 216 candidate SNPs significantly associated (p < 1e-5) with the trait. Although the use of CRISPR technology was suggested to understand how each of these SNPs would affect the phenotype, no functional validation was performed in this study (Katzenberger et al. 2015).

We applied FlyCADD to complement p-values in ranking the candidate SNPs based on predicted functional impact. Comparison of the GWAS p-values to the FlyCADD-scores of significantly associated SNPs did not reveal a significant correlation between the two metrics (Figure 9a). Applying FlyCADD scores showed that variants with significant p-values can even have a very low (< 0.2) predicted impact, and alternatively, variants with lower yet significant p-values can have high predicted impact (Figure 9a). For example, in the fifteen most significant candidate SNPs, nine variants have a predicted high impact above 0.6, and six variants have a predicted low (< 0.3) impact. Nine of these SNPs are in a 2000 bp intergenic region on chromosome 2L with two of these SNPs having low predicted impact and seven having high predicted impact (Figure 9b). This suggests the presence of both causal (potentially those with high FlyCADD scores) and hitchhiking (potentially those with low FlyCADD scores) SNPs among this panel of candidate SNPs with significant GWAS-derived p-values. Therefore, FlyCADD impact prediction scores apply an additional layer of information with regards to fitness-associated traits applicable to SNPs detected through GWAS.

**Figure 9:**
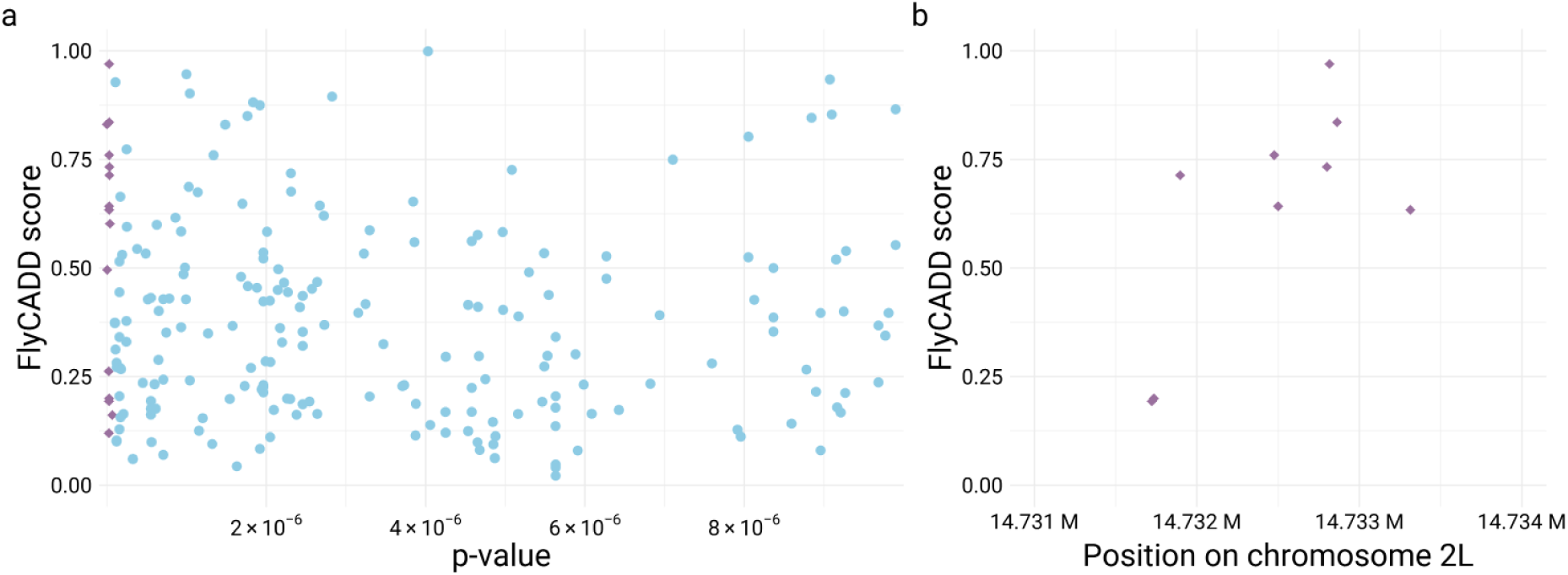
FlyCADD score and p-value compared for (a) 216 SNPs associated to death after brain trauma in GWAS by Katzenberger et al. (Katzenberger et al. 2015). The fifteen SNPs with the lowest p-values are highlighted in purple. (b) Nine of these SNPs with the lowest p-values are in a 2000 bp intergenic region of chromosome 2L and scores indicate predicted functional impact.

#### Ranking to identify causal SNP(s) or eliminate hitchhiking SNPs before experimental studies

Prioritization of SNPs using impact prediction scores helps eliminate hitchhiking loci and focus follow-up studies on functionally relevant variants, which will ultimately clarify the genetic architecture of complex traits. FlyCADD scores can be applied to either rank SNPs prior to functional studies or to confirm or interpret results of functional studies. To show how prioritization using FlyCADD scores can be applied to GWAS-identified SNPs and how this compares to experimental studies, we apply FlyCADD to the results of a GWAS on a fitness-affecting phenotype.

Natural variation in pigmentation has been observed among *D. melanogaster* populations, which can affect fitness by influencing temperature tolerance, mate choice, energy allocation and visibility among others (Dembeck et al. 2015; Freoa et al. 2023). Moreover, experimental evolution demonstrated that darker *D. melanogaster* populations showed a higher fecundity and significantly reduced lifespan compared to the less pigmented populations (Rajpurohit et al. 2016). Bastide et al. performed a GWAS identifying naturally occurring SNPs associated to female abdominal pigmentation, which resulted in seventeen significant candidate SNPs and hundreds of non-significant SNPs associated to a darker phenotype (Bastide et al. 2013). The three most significantly associated SNPs were located in the cis-regulatory region of *tan*, a pleiotropic gene involved in pigmentation, evolution of this trait, and other roles such as vision or mating behavior, suggesting that these SNPs affect pigmentation through regulation of *tan* (Gibert et al. 2017; Massey et al. 2019). Transgenic lines with each one out of eight combinations of these three SNPs were screened to determine the functional impact of the SNPs *in vivo* (Gibert et al. 2017). This demonstrated that all three SNPs (X:9227096, X:9227061, X:9226889) showed an additive, complex epistatic effect on pigmentation with most of the variance in pigmentation explained by X:9227096. In addition, the allele at X:9227061, associated with the dark phenotype in GWAS, resulted in less abdominal pigmentation in transgenic lines. This effect is in the opposite direction of what the GWAS initially suggested, which Gibert et al. attributed to linkage of the SNP with the other candidate SNPs as epistasis showed a less strong effect (Gibert et al. 2017).

We applied FlyCADD to complement the GWAS results and *in vivo* study by ranking these SNPs based on predicted functional impact. The FlyCADD impact prediction scores for the SNPs are as follows: 0.30 for X:9227096, 0.04 for X:9227061, 0.41 for X:9226889 (Figure 10). With FlyCADD scores between 0.30 and 0.60, two of these SNPs are predicted to have a moderate phenotypic impact. The low FlyCADD score of X:9227061 (0.04) suggests that its association to pigmentation in the GWAS was due to linkage with functionally relevant SNPs rather than a direct phenotypic effect. This aligns with the findings of Gibert et al. where functional assays on this the SNP showed opposite phenotypic effects compared to the GWAS prediction (Gibert et al. 2017). Based on FlyCADD scores, researchers would have been able to identify and potentially eliminate this SNP with little functional relevance prior to the functional validation study. The additional GWAS-identified SNPs, which have not yet been functionally validated (triangles in Figure 10) can now be ranked to focus future functional studies on those with highest FlyCADD scores, and, therefore, highest potential impact. The prioritization of candidate SNPs using FlyCADD scores facilitates targeted functional assays, focusing on top-ranked SNPs by filtering out (possibly linked) SNPs with lower functional impact scores such as X:9227061.

**Figure 10:**
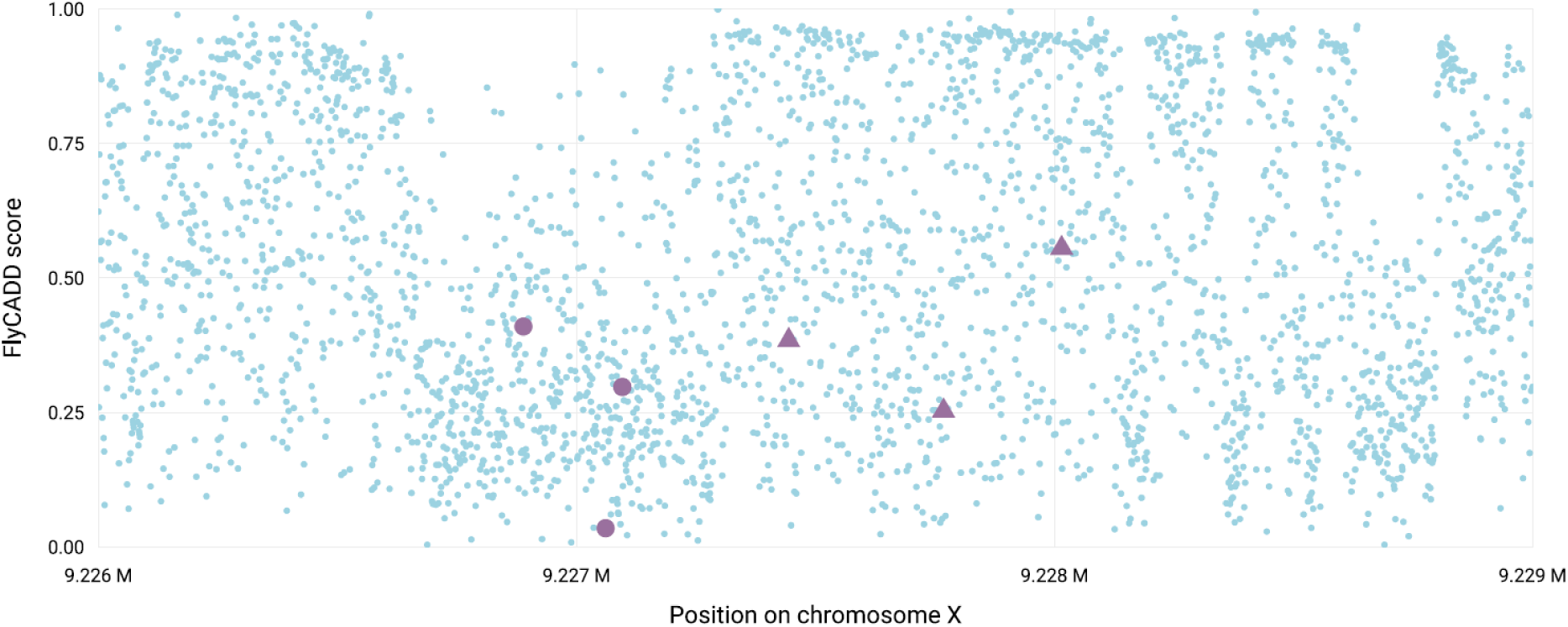
Mean FlyCADD scores per position (blue) for all possible single nucleotide variants in the regulatory domain associated with *tan*, with the FlyCADD score of the GWAS identified SNPs in purple of which the SNPs indicated with circles were functionally validated, and triangles were not functionally validated.

#### Evaluation of off-target effects in functional studies

Genome editing using CRISPR-Cas9 or other gene-editing technologies is widely used for functional genomics, both *in vivo* and in cell lines (Adli 2018). However, these techniques can introduce off-target effects, including unintended point mutations that may confound experimental results (Zhang et al. 2015). Additionally, intentional modifications in the target regions are required in some cases to increase efficiency of genome editing, for example by disrupting the protospacer adjacent motif (PAM) sites essential to Cas9 (Perlmutter et al. 2024; Zhang et al. 2015). These are not off-target effects of the CRISPR-Cas9 mechanism but are deliberate alterations to improve genome editing outcomes. FlyCADD can be applied in genome editing experiments to prioritize variants of interest, to evaluate the potential functional impact of intentional modifications essential in experimental design, as well as to evaluate the potential functional impact of unintended off-target effects resulting from genome editing. This can increase the confidence that observed phenotypic effects result from the on-target modification of interest, rather than from confounding (intentional or unintended) off-target point mutations.

In their study, Perlmutter et al. applied CRISPR-Cas9 to assess functional differences between two alleles of a non-synonymous polymorphism in the gene *Metchnikowin* (*Mtk*) in *D. melanogaster* (Perlmutter et al. 2024). *Mtk* is involved in microbe defense, response to wounding, neuropathology, and fitness among others (Levashina et al. 1998; Perlmutter et al. 2024; Swanson et al. 2020). Perlmutter et al. focused on infection and life-history assessments of CRISPR-Cas9 edited and control lines (Perlmutter et al. 2024). In addition to the desired single nucleotide modification, multiple silent point mutations were intentionally introduced in the target region to increase efficiency. The authors deemed the effect of these intended point mutations neglectable as these were synonymous mutations and limited expression differences were observed (Perlmutter et al. 2024).

We applied FlyCADD to determine whether the introduced silent mutations could have altered the results of the phenotypic screening. The SNP of interest (2R:15409024) has a predicted moderate functional impact with a FlyCADD score of 0.47, and the introduced silent mutations have FlyCADD scores of 0.52 (2R:15409016), 0.34 (2R:15409019) and 0.10 (2R:15409033).

These impact prediction scores reveal that, although the intentional point mutations are synonymous, they are predicted to have an impact on fitness, with one of these silent mutations (2R:15409016) having a predicted impact higher than the predicted impact of the SNP interest. These intended “silent” point mutations could potentially have an impact on the measured fitness-related phenotypes and therefore alter the phenotypic outcomes of the study. FlyCADD scores can now be applied during experimental design to more precisely place intended mutations and thereby avoid phenotypic effects caused by predetermined mutations.

Additionally, FlyCADD can be utilized after genome editing by applying the impact prediction scores to sequencing data of the obtained lines to estimate whether observed phenotypic differences are influenced by unintended off-target modifications. Perlmutter et al. identified 69 SNPs differing between the edited and unedited lines, most of which were deemed cases of mapping uncertainty (Perlmutter et al. 2024). Of these, however, 14 SNPs have a FlyCADD scores indicating a probability of being impactful (> 0.60), therefore it is valuable to further assess mapping quality for these SNPs to make sure these SNPs do not confound the phenotypic results observed. The authors identified one missense SNP (2R:16854417) as a confirmed off-target mutation segregating between the edited and unedited lines (Perlmutter et al. 2024). This SNP has a very high FlyCADD score (0.99), indicating that the SNP is very likely to impact fitness and potentially influenced the experimental results. This information on predicted variant impact allows for a more cautious interpretation of phenotypic measurements.

## Discussion

FlyCADD is a versatile new tool for genetics research in *D. melanogaster*, offering valuable insights into the functional impact of SNPs across the entire genome. By integrating 691 genomic features into a single impact prediction score, it enables researchers to prioritize SNPs, assess the potential phenotypic impact of natural variation, evaluate off-target effects of genome editing, and clarify patterns of causal and neutral variants based on predicted functional impact. The CADD framework is based on the assumption that (nearly) fixed derived variants are depleted of impactful mutations, whilst purifying selection has not affected simulated variants, and therefore, the set of simulated variants contains variants on the full spectrum of impact (Kircher et al. 2014). It is important to note that while high impact scores indicate functional impact, they do not necessarily imply a harmful or deleterious fitness effect. Variants with a high predicted impact score have a higher probability of having an impact on fitness, either harmful or beneficial. FlyCADD can advance our understanding of the functional importance of SNPs and genotype-phenotype connections in *D. melanogaster* by providing impact scores for every SNP throughout the genome.

### Ensuring model reliability through training and validation

The CADD framework works best when rich genomic information is available, such as in model species. The available sequencing datasets, high-quality reference genome, species alignments and nucleotide-based annotations for *D. melanogaster* made it feasible to develop FlyCADD tailored to *D. melanogaster* based on the existing CADD framework (Groß et al. 2020b; Hoskins et al. 2015; Kim et al. 2024; Kircher et al. 2014; Mackay et al. 2012a; Nunez et al. 2024; Rech et al. 2022). Ancestral sequence reconstruction was performed on a node that resulted in an ancestral sequence with minimized bias towards specific chromosomes or coding sequence, though challenges remained in reconstruction of heterochromatic regions such as the Y chromosome, repeat regions and centromeric regions as they are often lacking in genome assemblies (Chang and Larracuente 2018). Therefore, FlyCADD is trained on the autosomes and X chromosome, and no scores are present for variants on the Y chromosome. The reconstructed ancestral genome was sufficiently large to develop a robust predictive model trained on more than five million annotated single nucleotide variants. Our data shows that the CADD approach can be effectively extended to insect genomes, especially when high-quality genomic resources are present.

Our analyses indicate that FlyCADD has a high accuracy with ROC-AUC of 0.83 and a prediction accuracy of 0.76 on the held-out test dataset. These numbers indicate that FlyCADD performance is higher compared to the existing mouse (ROC-AUC of 0.67) and pig (ROC-AUC of 0.68) CADD models (Groß et al. 2018; Groß et al. 2020b). mCADD (mouse) and humanCADD (hCADD, (Rentzsch et al. 2018; Schubach et al. 2024)), tested on available validation datasets containing variants of known pathogenicity rather than held-out training data, reported AUC scores of > 0.95. In contrast, there is no extensive, genome-wide validation dataset available for functional impact of SNPs in *D. melanogaster* beyond the binary phenotypic class of lethal point mutations from FlyBase included in the current study for score validation. Therefore, direct comparison of model accuracy is difficult as the CADD models are species-specific. However, validation of FlyCADD scores using known lethal mutations demonstrated that FlyCADD can correctly predict functional impact for experimentally tested point mutations and identify the causal mutation when multiple point mutations co-occur. This is consistent with findings from other variant effect predictors, such as ProteoCast, which also identify the impact of these mutations and highlight the challenge of disentangling co-occurring genetic variants (Abakarova et al. 2025).

### Species-specific functional predictions

Species-specific models are important as, for example, mammal and insect genomes differ widely, and therefore, predictability of functional impact of SNPs and importance of different genomic features differ between species (Huber et al. 2017). Previous CADD models demonstrated that annotations derived from the sequence contribute most to the predictions and that species-specific annotations are enhancing performance (Groß et al. 2018). Out of a total of 691 features in FlyCADD, the most important features were derived from the sequence and were combinations of positional, functional or conservation scores, emphasizing the importance of composite features over single annotations for *D. melanogaster*. Species-specific application of the CADD framework is important to preserve the differences between species and accurately predict functional impact for the species of interest. As a species-specific implementation of the CADD framework, FlyCADD enhances genotype–phenotype research in *D. melanogaster* by providing predictions optimized for the genomic architecture of this species.

It is currently unknown whether species-specific CADD models can reliably be applied to (closely) related species, for example applying FlyCADD to *D. suzukii*, *D. simulans* or *D. yakuba*. ChickenCADD (chCADD, (Groß et al. 2020a)) has been applied to the pink pigeon genome, but no validation was performed to assess the accuracy of chCADD on this pink pigeon genome (Speak et al. 2024). The availability of genome editing techniques and high-quality genomic resources in *D. melanogaster* and related *Drosophila* species provides opportunities to functionally validate impact prediction scores from FlyCADD within and among related species. Future research could evaluate FlyCADD’s transferability to other *Drosophila* species through score lift over or by directly scoring novel SNPs, as the scripts and impact prediction scores of FlyCADD are publicly available.

### Genome-wide functional insights from FlyCADD

CADD applied to the human genome is widely used for analysis of disease-related polymorphisms, while pCADD and chCADD have been applied to agriculture and conservation genomics (Derks et al. 2021; Groß et al. 2020b; Speak et al. 2024). By applying FlyCADD to different use cases, we demonstrated several examples of its value in prioritizing SNPs from GWAS prior to follow-up studies, design and evaluation of genome editing experiments and assessment of natural variation in the fruit fly. FlyCADD enables researchers to examine functional impact of candidate loci associated to complex traits at the SNP level across the entire *D. melanogaster* genome and in different research contexts, offering insights into the genetic basis of phenotypic variation. Given that linkage between causal SNPs and hitchhiking loci is common, both in natural and experimental populations, FlyCADD can help distinguish functionally impactful SNPs from neutral loci based on combined annotations, especially in non-coding regions where traditional methods of assessing functional impact of SNPs fall short (Smit-McBride et al. 1988; Smith and Haigh 1974). We have shown that FlyCADD scores can easily be applied as an additional layer of information on candidate SNPs and as scores are available across the entire genome, no variant or region is overlooked. Currently, FlyCADD can only be used to score functional impact of point mutations and not structural variants or other types of genomic variants. Future expansion of FlyCADD towards predicting functional impact of other types of genomic variants is possible, however. For example, the CADD framework has been applied to score functional impact of structural variants in the human genome (Kleinert and Kircher 2022).

In conclusion, we present FlyCADD, an impact prediction tool for SNPs in the genome of *D. melanogaster.* Besides the selected use cases, FlyCADD impact prediction scores can be used on a genome-wide scale in many more research contexts to study the genetic basis of phenotypes in *D. melanogaster*. Prioritization of promising SNPs or elimination of predicted low-impact SNPs based on FlyCADD impact prediction scores will help researchers reduce time, cost, and effort of follow-up studies by preventing unnecessary validation of hitchhiking loci that are not likely to impact the phenotype of interest. This targeted approach directs resources toward functionally relevant SNPs, making (experimental or computational) validation of impact more feasible.

We have made FlyCADD impact prediction scores readily available on Zenodo, providing both precomputed scores for all possible single nucleotide variants on the *D. melanogaster* reference genome and a locally executable pipeline for scoring novel variants of interest (https://doi.org/10.5281/zenodo.14887337). By extending functional SNP prediction beyond coding regions and known functional domains, FlyCADD presents a valuable addition to the existing genetic toolbox of *Drosophila*. The combination of FlyCADD with the powerful functional genetic tools for *D. melanogaster* provides unique opportunities to create a better understanding of the genotype-phenotype connections underlying phenotypic variation.

## Data availability

File S1 describes the Cactus 166-way multi-species alignment that was used for ancestral sequence reconstruction and details on the extracted ancestral sequence, including two alternative ancestral sequences that showed excessive biases. File S2 gives an overview of the annotations and combinations thereof used by FlyCADD. File S3 shows the weight of all features incorporated in the trained logistic regression model, describing the contribution of each feature to the impact prediction.

All supporting data and resources related to FlyCADD are publicly available. The pipeline of FlyCADD development is available at GitHub (https://github.com/JuliaBeets/FlyCADD). Precomputed scores, a locally executable FlyCADD pipeline (including the trained logistic regression model files, annotation files and scripts) for scoring novel variants, the multi-species alignment file, the reconstructed ancestral sequence, generated derived and simulated variants, and FlyCADD scores at all codon positions of unique transcripts in the *D. melanogaster* genome can be found on Zenodo (https://doi.org/10.5281/zenodo.14887337).

## Acknowledgements

The authors acknowledge BAZIS HPC cluster computing facilities at the Vrije Universiteit Amsterdam. The authors thank the *D. melanogaster* research community for their contributions in generating essential resources, annotations, and sequencing datasets that formed the basis for developing FlyCADD. We acknowledge the *Drosophila* Evolutionary Population Genomics Consortium (DrosEU) for generating and maintaining essential sequencing datasets and genomic resources, including the DEST(2.0) resource.

The authors have declared no competing interest.

