## Supplementary File S1 for "Predicting the functional impact of single nucleotide variants in *Drosophila melanogaster* with FlyCADD"

**File S1: Ancestral sequence reconstruction**

The FlyCADD development is based on comparisons between the *D. melanogaster* reference genome and a reconstructed ancestral genome. The phylogenetic tree of the alignment that was used is shown in Figure 1 with *D. melanogaster* in purple. This multi-species alignment is a 166-way alignment made available for FlyCADD development, whereas a 298-way alignment, including the 166 species subset, is published (Kim et al. 2024). The alignment file of the 166-way alignment specific to this study is available on Zenodo (10.5281/zenodo.14887338).

Three ancestral sequences at nodes A, B and C were assessed for ancestral sequence quality, overlap with coding sequence (CDS) and potential bias towards specific chromosomes. The ancestral reconstruction of node B, encompassing the branches in green, was picked. Table 1 describes the size of the ancestral sequence reconstruction, its distribution across the different chromosomes, and the percentage of the reference genome coding sequence covered by the ancestral sequence. This is depicted in Figure 2 by showing the distribution of coding sequence on the reference genome (purple) and the distribution of reconstructed ancestral sequence along the reference genome (orange).

The aim was to choose a reconstructed ancestral sequence that fit three criteria: the largest reconstructed ancestral sequence to maximize the training dataset, the least chromosome bias, and the least coding sequence bias. To assess potential bias of the reconstructed ancestral sequence towards coding regions, we compared the overall percentage of coding sequence in the *D. melanogaster* genome to the percentage of overlap between coding sequence and the respective reconstructed ancestral sequence. The percentage of CDS across the *D. melanogaster* reference genome is ~ 15%. This indicates that an ancestral sequence closer to the species of interest (Node C) shows ancestral sequence reconstruction bias towards excessive coding sequence coverage. Node C also resulted in an ancestral sequence with reconstruction bias towards chromosomes 3 and X, and away from chromosomes 2R (harbouring only 2.3 % of the ancestral sequence) and 4 (harbouring only 0.01 % of the ancestral sequence). Node B yielded the largest reconstructed ancestral sequence (covering 16.4 % of the reference genome), while the furthest node (A) reduces the overall percentage of reconstructed ancestral sequence, ultimately resulting in too few variants for model training. The overall percentage of ancestral sequence overlapping with CDS was comparable between nodes A (17.77 %) and B (18.47 %). Therefore, we picked the reconstructed ancestral sequence of node B for FlyCADD development, which showed the least CDS or chromosome biases and is largest by covering ~ 16 % of the reference genome.

**Figure 1:** Species tree of the 166-way alignment used to infer the ancestral sequence of the modern *D. melanogaster* in purple. From the assessed nodes (blue), the reconstructed ancestral sequence of node B was chosen for FlyCADD.

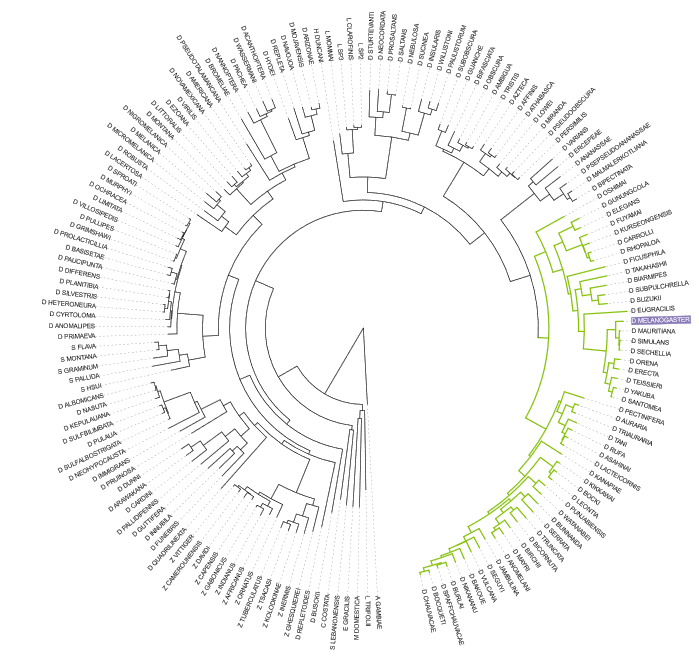

A

B	C

| **Table 1:** Percentage of genome reconstructed at three nodes from the alignment. Additionally, for each chromosome, the left column shows the proportion of the reconstructed ancestral sequence at that chromosome, reflecting the ancestral sequence distribution. The right column shows the percentage of reconstructed ancestral sequence overlapping with coding sequence (CDS) of this chromosome, as measure for potential reconstruction bias toward coding regions. The final row indicates the overall percentage of genome-wide overlap between the reconstructed ancestor and the reference genome CDS. The reconstructed ancestral sequence from node B was used for FlyCADD development. | | | | | | |
| --- | --- | --- | --- | --- | --- | --- |
|  | **Node A** | | **Node B** | | **Node C** | |
| **% reconstructed** | 10.4 | | 16.4 | | 13.5 | |
| **2L** | 17.9 | 30.69 | 21.3 | 29.42 | 15.6 | 31.33 |
| **2R** | 18.8 | 0.69 | 16.9 | 0.69 | 2.3 | 0.38 |
| **3L** | 21.9 | 26.15 | 21.7 | 26.38 | 28.5 | 22.79 |
| **3R** | 25.1 | 2.53 | 24.3 | 4.38 | 33.7 | 21.36 |
| **4** | 0.5 | 17.04 | 0.7 | 19.27 | 0.01 | 57.64 |
| **X** | 15.8 | 36.20 | 15.1 | 34.15 | 18.8 | 23.99 |
| **% overlap with CDS** | 17.77 | | 18.47 | | 23.34 | |

**Figure 2:** Distribution of ancestral sequence (orange) across the autosomes and X chromosome of *D. melanogaster* for reconstructed ancestors at three different nodes a) node A, b) node B and c) node C (see File S1 Figure 1). The distribution of coding sequence along the reference genome is depicted in purple. Darker colours indicate higher density of coding sequence (purple) or reconstructed ancestral sequence (orange).

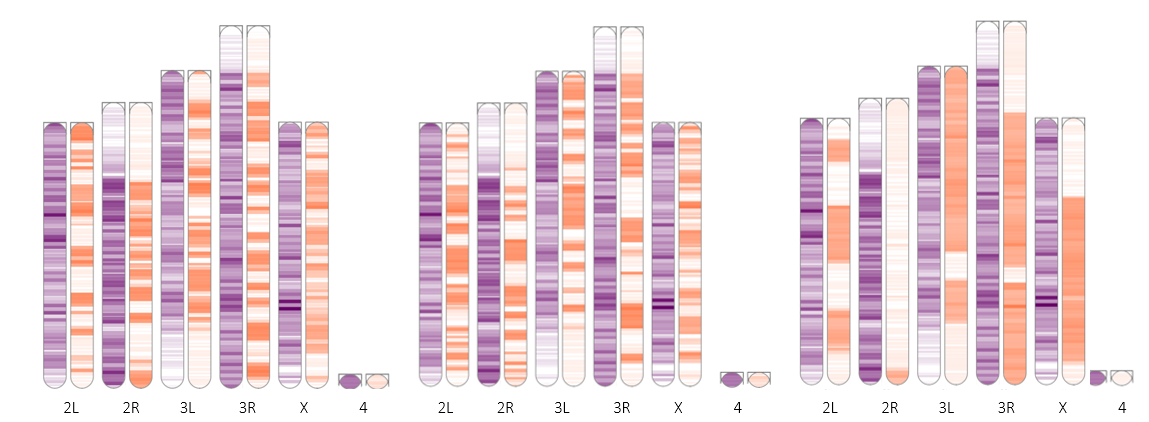

a	b	c

**References File S1**

Kim BY, Gellert HR, Church SH, Suvorov A, Anderson SS, Barmina O, Beskid SG, Comeault AA, Crown KN, Diamond SE et al. 2024. Single-fly genome assemblies fill major phylogenomic gaps across the drosophilidae tree of life. PLOS Biology. 22(7):e3002697. doi:10.1371/journal.pbio.3002697.
