## Supplementary File S2 for "Predicting the functional impact of single nucleotide variants in *Drosophila melanogaster* with FlyCADD"

**File S2: Overview of the annotations in FlyCADD**

File S2 Table 1 describes the individual annotations used for training, testing and application of FlyCADD. Combinations between annotations were made for the annotations in File S2 Table 2. The total number of features in FlyCADD was 691, descriptions per annotation can be found in File S1.

| **Table 1:** Individual annotations used in development and application of FlyCADD. From each annotation, the name in the model, description, datatype and source are described. Annotations with multiple possible categories such as Consequence, RepeatType and amino acids were divided into separate features. | | | |
| --- | --- | --- | --- |
| **Annotation** | **Description** | **Datatype** | **Source** |
| #Derived(1) _Simulated(0)* | Variant belongs to derived or simulated variant set | Boolean | FlyCADD development |
| Chrom** | Chromosome | Categorical | Release 6 reference genome |
| Pos** | Genomic position (1-based) | Integer | Release 6 reference genome |
| Ref | Reference allele | Categorical | Release 6 reference genome |
| Alt | Alternative allele | Categorical | FlyCADD development |
| isTv | Transversion | Boolean | Ensembl Variant Effect Predictor (McLaren et al. 2016) |
| Consequence | Functional consequence (SG stop-gain, IF inframe-insertion, SL stop-loss, S splice-site, SN synonymous, DN downstream, R regulatory region, U3 3Prime_UTR, O unknown, NS non-synonymous, FS frameshift, CS canonical splice, NC non-coding change, IG intergenic, UP upstream, U5 5Prime-UTR, I intronic) | Categorical | Ensembl Variant Effect Predictor |
| GC | GC content percentage in +/- 75 bp window | Numerical | Ensembl Variant Effect Predictor |
| CpG | CpG percentage in +/- 75 bp window | Numerical | Ensembl Variant Effect Predictor |
| Domain | Domain (lcompl, ncoils, ndomain, sigp, tmhmm, -) | Categorical | Ensembl Variant Effect Predictor |
| oAA | Amino acid with reference allele | Categorical | Ensembl Variant Effect Predictor |
| nAA | Amino acid with alternative allele | Categorical | Ensembl Variant Effect Predictor |
| Grantham | Grantham score for amino acid change | Integer | Grantham matrix (Grantham 1974) |
| cDNApos | Proximity to transcription start site | Integer | Ensembl Variant Effect Predictor |
| relcDNApos | Relative position in the coding region | Numerical | Ensembl Variant Effect Predictor |
| CDSpos | Proximity to start of coding sequence | Integer | Ensembl Variant Effect Predictor |
| relCDSpos | Relative position in the coding region | Numerical | Ensembl Variant Effect Predictor |
| protPos | Position of the amino acid in the protein | Numerical | Ensembl Variant Effect Predictor |
| relProtPos | Relative position of the amino acid in the protein | Integer | Ensembl Variant Effect Predictor |
| PhastCons | PhastCons conservation score based on 27-way alignment | Numerical | UCSC (Perez et al. 2025) |
| PhyloP | PhyloP conservation score based on 27-way alignment | Numerical | UCSC (Perez et al. 2025) |
| Roll | Predicted local DNA shape feature Roll | Numerical | DNAshapeR (Chiu et al. 2016) |
| EP | Predicted electrostatic potential of local DNA | Numerical | DNAshapeR (Chiu et al. 2016) |
| MGW | Predicted local DNA shape feature minor groove width | Numerical | DNAshapeR (Chiu et al. 2016) |
| HelT | Predicted local DNA shape feature helix twist | Numerical | DNAshapeR (Chiu et al. 2016) |
| ProT | Predicted local DNA shape feature propeller twist | Numerical | DNAshapeR (Chiu et al. 2016) |
| Repeats | Repeat | Boolean | UCSC (Perez et al. 2025) |
| Isgene | Coding region | Boolean | UCSC (Perez et al. 2025) |
| RepeatType | Repeat type (LINE, LTR, DNA, Simple, Low Complexity, Satellite, RNA, Unknown, No repeat) | Categorical | UCSC (Perez et al. 2025), RepeatMasker track |
| TFBS | Transcription factor binding site | Boolean | RedFly (Keränen et al. 2022) |
| miRNA | microRNA-encoding sequence | Boolean | FlyBase 6.54 (Öztürk-Çolak et al. 2024) |
| ReMap | Density of ReMap records for regulatory elements | Numerical | UCSC (Perez et al. 2025) |
| CRM | (predicted) cis-regulatory motif | Boolean | RedFly (Keränen et al. 2022) |
| PhastCons124 | PhastCons conservation score based on 124-way alignment | Numerical | UCSC (Perez et al. 2025) |
| PhyloP124 | PhyloP conservation score based on 124-way alignment | Numerical | UCSC (Perez et al. 2025) |
| BG3_state | Chromatin state in BG3 cells | Categorical | FlyBase 6.54 (Öztürk-Çolak et al. 2024) |
| S2_state | Chromatin state in S2 cells | Categorical | FlyBase 6.54 (Öztürk-Çolak et al. 2024) |
| PhyloP | PhyloP conservation scores based on Cactus alignment | Numerical | Petrov lab (Kim et al. 2024) |
| GERP-RS | Rejected substitution score GERP based on Cactus alignment | Numerical | Petrov lab (Kim et al. 2024) |
| GERP-N | Neutral rate of evolution score GERP based on Cactus alignment | Numerical | Petrov lab (Kim et al. 2024) |

*Only applied in training and testing.

**Not applied for training, testing or computing scores.

| **Table 2:** Combined features of annotations. In each row, every feature from selection 1 is paired with every feature from selection 2 | |
| --- | --- |
| **Selection 1** | **Selection 2** |
| Ref (A, C, T, G) | Alt (A, C, T, G) |
| oAA (A, R, N, D, C, E, Q, G, H, I, L, K, M, F, P, S, T, W, Y, V, *) | nAA (A, R, N, D, C, E, Q, G, H, I, L, K, M, F, P, S, T, W, Y, V, *) |
| VEP functional consequence (SG stop-gain, IF inframe-insertion, SL stop-loss, S splice-site, SN synonymous, DN downstream, R regulatory region, U3 3Prime_UTR, O unknown, NS non-synonymous, FS frameshift, CS canonical splice, NC non-coding change, IG intergenic, UP upstream, U5 5Prime-UTR, I intronic) | cDNApos, CDSpos, PhastCons, PhyloP, protPos, relcDNApos, relCDSpos, relprotPos, PhastCons124, PhyloP124, phylop_all, ReMap, GERPS_all, GERPN_all |

**References File S2**

Chiu TP, Comoglio F, Zhou T, Yang L, Paro R, Rohs R. 2016. Dnashaper: An r/bioconductor package for DNA shape prediction and feature encoding. Bioinformatics. 32(8):1211-1213. doi:10.1093/bioinformatics/btv735.

Grantham R. 1974. Amino acid difference formula to help explain protein evolution. Science. 185(4154):862-864. doi:10.1126/science.185.4154.862.

Keränen SVE, Villahoz-Baleta A, Bruno AE, Halfon MS. 2022. Redfly: An integrated knowledgebase for insect regulatory genomics. Insects. 13(7):618. doi:10.3390/insects13070618.

Kim BY, Gellert HR, Church SH, Suvorov A, Anderson SS, Barmina O, Beskid SG, Comeault AA, Crown KN, Diamond SE et al. 2024. Single-fly genome assemblies fill major phylogenomic gaps across the drosophilidae tree of life. PLOS Biology. 22(7):e3002697. doi:10.1371/journal.pbio.3002697.

McLaren W, Gil L, Hunt SE, Riat HS, Ritchie GRS, Thormann A, Flicek P, Cunningham F. 2016. The ensembl variant effect predictor. Genome Biology. 17(1):122. doi:10.1186/s13059-016-0974-4.

Öztürk-Çolak A, Marygold SJ, Antonazzo G, Attrill H, Goutte-Gattat D, Jenkins VK, Matthews BB, Millburn G, dos Santos G, Tabone CJ, FlyBase C. 2024. Flybase: Updates to the drosophila genes and genomes database. Genetics. 227(1):iyad211. doi:10.1093/genetics/iyad211.

Perez G, Barber GP, Benet-Pages A, Casper J, Clawson H, Diekhans M, Fischer C, Gonzalez JN, Hinrichs AS, Lee CM et al. 2025. The ucsc genome browser database: 2025 update. Nucleic Acids Res. 53(D1):D1243-d1249. doi:10.1093/nar/gkae974.
