## Supplementary File S3 for "Predicting the functional impact of single nucleotide variants in *Drosophila melanogaster* with FlyCADD"

**File S3: Contribution of all 691 features in FlyCADD**

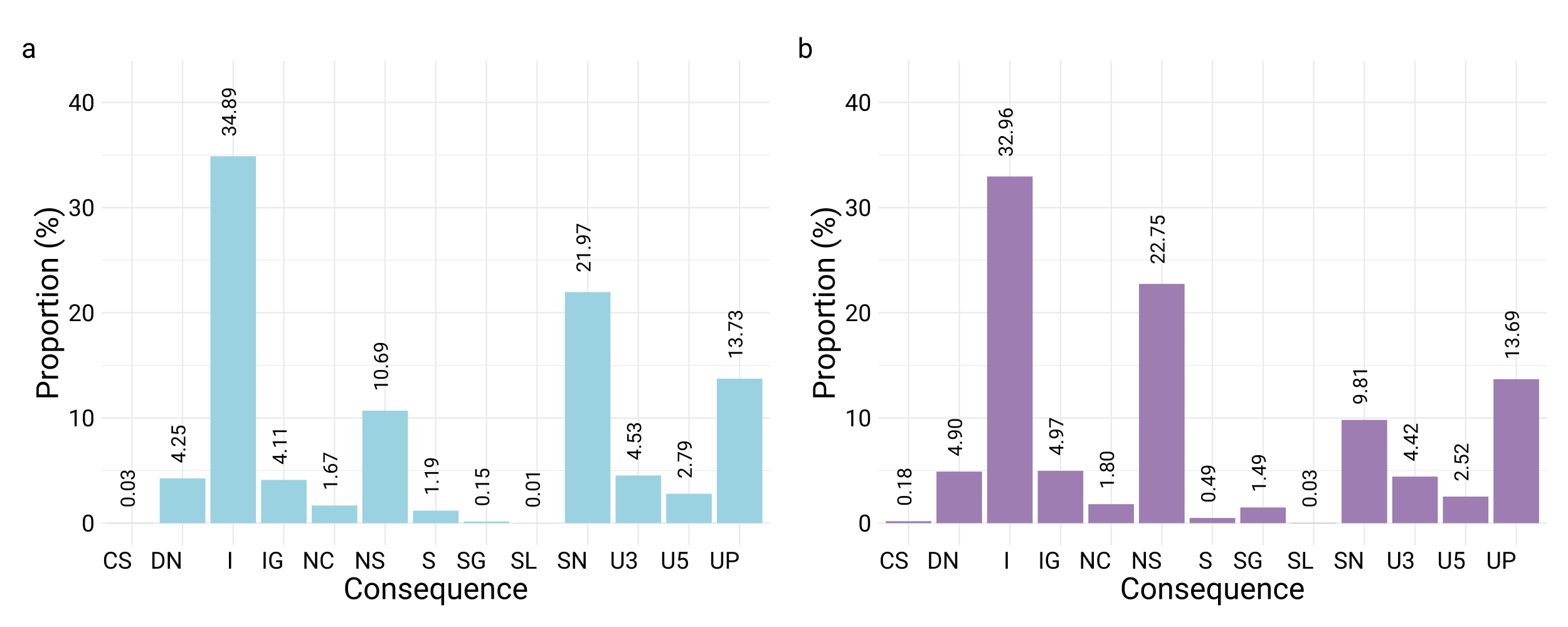
**Proportion of variants per consequence present in training data**

**Figure 1**: Proportion of variants present in the (a) derived variant set and (b) simulated variant set with different consequences. CS: canonical splice; DN: downstream; I: intronic; IG: intergenic; NC: non-coding exon change; NS: non-synonymous; S: splice site; SG: stop gain; SL: stop loss; SN: synonymous; U3: 3 prime UTR; U5: 5 prime UTR; UP: upstream.

**337 features with negative weight**

| **Table 1:** Features with negative weight. | | | | | | | |
| --- | --- | --- | --- | --- | --- | --- | --- |
| **Feature** | **Weight** |  | **Feature** | **Weight** |  | **Feature** | **Weight** |
| Consequence_SG | -6.69315 |  | Ref_C | -0.12363 |  | DN_GERPRS_all | -0.05730 |
| SL_relCDSpos | -2.66447 |  | IG_PhyloP124 | -0.11934 |  | UP_GERPRS_all | -0.05394 |
| SN_CDSpos | -0.85041 |  | Ref_G | -0.11853 |  | T_T | -0.05321 |
| CDSpos | -0.75054 |  | nAA_S | -0.11222 |  | IND_CDSpos | -0.05070 |
| nAA_* | -0.71096 |  | DN_PhyloP124 | -0.11098 |  | IND_protPos | -0.05070 |
| NS_relCDSpos | -0.61073 |  | L_M | -0.10658 |  | oAA_UD | -0.05070 |
| PhyloP | -0.57155 |  | U3_PhyloP | -0.10595 |  | nAA_UD | -0.05070 |
| (intercept) | -0.39964 |  | I_GERPRS_all | -0.10276 |  | U3_PhyloP124 | -0.05069 |
| SN_relprotPos | -0.37074 |  | SG_PhastCons | -0.10075 |  | A_A | -0.05053 |
| NS_PhyloP | -0.35376 |  | IND_Grantham | -0.10046 |  | NS_cDNApos | -0.04994 |
| L_L | -0.29314 |  | SN_phylop_all | -0.09567 |  | Y_* | -0.04976 |
| E_K | -0.28131 |  | L_I | -0.09530 |  | IG_GERPRS_all | -0.04947 |
| nAA_L | -0.27964 |  | nAA_P | -0.09407 |  | U3_GERPRS_all | -0.04827 |
| I_PhyloP | -0.27417 |  | L_R | -0.09336 |  | nAA_A | -0.04811 |
| E_D | -0.25255 |  | L_Q | -0.08777 |  | nAA_F | -0.04629 |
| E_E | -0.22621 |  | L_S | -0.08626 |  | SL_PhyloP | -0.04583 |
| nAA_C | -0.22094 |  | CS_GERPRS_all | -0.08603 |  | PhastCons124 | -0.04572 |
| S_relprotPos | -0.21909 |  | SN_GERPN_all | -0.08398 |  | NC_PhyloP124 | -0.04512 |
| S_CDSpos | -0.20027 |  | C.1_G.1 | -0.08377 |  | C_* | -0.04384 |
| E_* | -0.19843 |  | NS_PhastCons | -0.08276 |  | R_G | -0.04332 |
| CS_relCDSpos | -0.19834 |  | NC_PhyloP | -0.08041 |  | U5_PhyloP124 | -0.04328 |
| E_Q | -0.18889 |  | G.1_C.1 | -0.07918 |  | C.1_A.1 | -0.04184 |
| E_G | -0.18789 |  | SG_GERPRS_all | -0.07784 |  | G.1_T.1 | -0.04182 |
| I_PhyloP124 | -0.18723 |  | SG_GERPN_all | -0.07593 |  | R_* | -0.04118 |
| nAA_R | -0.18554 |  | nAA_D | -0.07473 |  | S_P | -0.03886 |
| UP_PhyloP | -0.18348 |  | PhyloP124 | -0.07322 |  | NS_GERPN_all | -0.03853 |
| nAA_E | -0.18188 |  | nAA_M | -0.07087 |  | nAA_T | -0.03736 |
| NS_CDSpos | -0.17909 |  | nAA_Q | -0.07046 |  | Q_H | -0.03605 |
| SL_relprotPos | -0.17883 |  | U5_PhyloP | -0.07034 |  | S_phylop_all | -0.03449 |
| UP_PhyloP124 | -0.17219 |  | SG_protPos | -0.06971 |  | cDNApos | -0.03087 |
| L_P | -0.16732 |  | SG_CDSpos | -0.06947 |  | GC | -0.03048 |
| E_V | -0.16457 |  | C.1_T.1 | -0.06940 |  | R_S | -0.02863 |
| GERPRS_all | -0.16399 |  | R_H | -0.06818 |  | P_P | -0.02837 |
| L_F | -0.15052 |  | PhastCons | -0.06789 |  | D_H | -0.02814 |
| L_V | -0.14279 |  | L_H | -0.06628 |  | S_W | -0.02681 |
| nAA_Y | -0.14134 |  | G.1_A.1 | -0.06597 |  | nAA_I | -0.02675 |
| nAA_K | -0.13761 |  | I_phylop_all | -0.06249 |  | F_S | -0.02609 |
| nAA_G | -0.13388 |  | R_P | -0.06229 |  | N_K | -0.02594 |
| E_A | -0.13343 |  | NS_PhastCons124 | -0.06199 |  | T_M | -0.02579 |
| DN_PhyloP | -0.13221 |  | K_N | -0.06144 |  | H_Q | -0.02563 |
| IG_PhyloP | -0.13179 |  | L_* | -0.06112 |  | U5_phylop_all | -0.02477 |
| CpG | -0.13073 |  | SN_PhyloP | -0.06060 |  | I_PhastCons | -0.02451 |
| NS_GERPRS_all | -0.12925 |  | nAA_V | -0.05980 |  | I_M | -0.02347 |
| nAA_W | -0.12864 |  | nAA_H | -0.05822 |  | G_R | -0.02327 |

| **Feature** | **Weight** |  | **Feature** | **Weight** |  | **Feature** | **Weight** |
| --- | --- | --- | --- | --- | --- | --- | --- |
| R_C | -0.02266 |  | V_D | -0.01000 |  | E_S | -0.00608 |
| S_* | -0.02255 |  | F_P | -0.00989 |  | oAA_A | -0.00586 |
| SL_PhastCons124 | -0.02152 |  | E_R | -0.00984 |  | L_A | -0.00566 |
| C_R | -0.02128 |  | S_PhyloP | -0.00983 |  | F_N | -0.00554 |
| V_F | -0.02122 |  | D_P | -0.00981 |  | M_A | -0.00547 |
| Y_N | -0.02115 |  | CS_CDSpos | -0.00963 |  | Domain_ncoils | -0.00545 |
| K_* | -0.02113 |  | CS_PhastCons | -0.00949 |  | M_P | -0.00543 |
| N_I | -0.02105 |  | G_P | -0.00938 |  | E_N | -0.00543 |
| NC_GERPRS_all | -0.02044 |  | M_T | -0.00928 |  | Y_S | -0.00542 |
| D_Y | -0.02018 |  | S_F | -0.00919 |  | A_Y | -0.00534 |
| R_Q | -0.01993 |  | I_PhastCons124 | -0.00904 |  | U5_PhastCons | -0.00519 |
| U3_phylop_all | -0.01956 |  | Y_H | -0.00899 |  | N_F | -0.00518 |
| phylop_all | -0.01941 |  | U3_PhastCons124 | -0.00895 |  | *_S | -0.00501 |
| Alt_T | -0.01929 |  | CRM | -0.00854 |  | K_Y | -0.00492 |
| D_V | -0.01919 |  | nAA_N | -0.00843 |  | IG_PhastCons124 | -0.00485 |
| SG_cDNApos | -0.01841 |  | W_G | -0.00827 |  | H_C | -0.00471 |
| L_W | -0.01831 |  | D_F | -0.00811 |  | K_T | -0.00469 |
| Alt_A | -0.01806 |  | Y_P | -0.00798 |  | C_F | -0.00464 |
| R_L | -0.01712 |  | C_W | -0.00796 |  | G_Q | -0.00463 |
| oAA_T | -0.01705 |  | NC_PhastCons124 | -0.00792 |  | I_W | -0.00463 |
| S_I | -0.01687 |  | K_H | -0.00788 |  | Y_E | -0.00454 |
| K_I | -0.01650 |  | I_S | -0.00771 |  | M_E | -0.00452 |
| T_R | -0.01648 |  | H_V | -0.00767 |  | M_* | -0.00449 |
| S2_state | -0.01620 |  | E_F | -0.00739 |  | Domain_lcompl | -0.00447 |
| W_R | -0.01614 |  | P_W | -0.00739 |  | RepeatType_LCR | -0.00444 |
| E_L | -0.01578 |  | W_P | -0.00738 |  | N_W | -0.00443 |
| I_F | -0.01455 |  | T_F | -0.00735 |  | M_M | -0.00442 |
| SL_cDNApos | -0.01402 |  | S_L | -0.00732 |  | I_T | -0.00437 |
| D_N | -0.01326 |  | *_Y | -0.00702 |  | S_ReMap | -0.00436 |
| I_N | -0.01305 |  | Q_M | -0.00697 |  | *_L | -0.00424 |
| NC_phylop_all | -0.01270 |  | N_M | -0.00695 |  | V_M | -0.00424 |
| SG_PhyloP124 | -0.01234 |  | V_G | -0.00682 |  | F_V | -0.00384 |
| G_* | -0.01204 |  | M_R | -0.00679 |  | SG_phylop_all | -0.00378 |
| K_D | -0.01162 |  | S_R | -0.00668 |  | W_H | -0.00376 |
| R_W | -0.01153 |  | Roll | -0.00667 |  | *_K | -0.00336 |
| K_G | -0.01149 |  | RepeatType_rRNA | -0.00654 |  | E_P | -0.00335 |
| *_Q | -0.01147 |  | *_R | -0.00654 |  | Y_V | -0.00326 |
| U5_GERPRS_all | -0.01140 |  | Y_G | -0.00632 |  | CS_ReMap | -0.00326 |
| RepeatType_  Simple_repeat | -0.01080 |  | P_* | -0.00629 |  | Domain_sigp | -0.00321 |
| Y_D | -0.01076 |  | CS_PhyloP | -0.00629 |  | S_PhyloP124 | -0.00307 |
| Q_* | -0.01075 |  | F_Q | -0.00627 |  | Y_I | -0.00298 |
| SL_phylop_all | -0.01074 |  | K_L | -0.00616 |  | L_Y | -0.00290 |
| SN_ReMap | -0.01064 |  | M_F | -0.00612 |  | SG_relcDNApos | -0.00269 |
| U3_relcDNApos | -0.01055 |  | L_T | -0.00610 |  | F_I | -0.00266 |
| A_T | -0.01021 |  | F_W | -0.00609 |  | L_C | -0.00263 |
| T_W | -0.01009 |  | Q_C | -0.00608 |  | E_Y | -0.00260 |

| **Feature** | **Weight** |  | **Feature** | **Weight** |  | **Feature** | **Weight** |
| --- | --- | --- | --- | --- | --- | --- | --- |
| F_* | -0.00260 |  | K_S | -0.00114 |  | N_P | -0.00029 |
| I_P | -0.00253 |  | U3_ReMap | -0.00111 |  | *_I | -0.00017 |
| NC_relcDNApos | -0.00248 |  | L_K | -0.00111 |  | oAA_I | -0.00017 |
| CS_cDNApos | -0.00247 |  | T_P | -0.00109 |  | F_A | -0.00016 |
| Q_R | -0.00246 |  | SN_relcDNApos | -0.00103 |  | Q_A | -0.00011 |
| R_I | -0.00244 |  | CS_PhastCons124 | -0.00101 |  | *_F | -8.69E-05 |
| Q_T | -0.00236 |  | I_R | -0.00098 |  | D_I | -8.41E-05 |
| U5_PhastCons124 | -0.00222 |  | G_N | -0.00095 |  | R_N | -6.04E-05 |
| L_D | -0.00219 |  | V_S | -0.00091 |  | A_I | -3.85E-05 |
| K_M | -0.00206 |  | D_R | -0.00089 |  | P_R | -2.16E-05 |
| L_G | -0.00189 |  | S_GERPN_all | -0.00087 |  | N_G | -8.28E-06 |
| E_C | -0.00184 |  | K_V | -0.00082 |  | I_ReMap | -5.90E-06 |
| Y_T | -0.00180 |  | Domain_tmhmm | -0.00081 |  | I_Q | -1.98E-07 |
| U5_cDNApos | -0.00170 |  | Q_S | -0.00081 |  | F_M | -1.83E-07 |
| V_H | -0.00164 |  | G_H | -0.00067 |  | G_Y | -1.83E-07 |
| DN_ReMap | -0.00151 |  | V_* | -0.00062 |  | H_W | -1.45E-07 |
| F_T | -0.00147 |  | F_R | -0.00053 |  | C_I | -1.32E-07 |
| I_I | -0.00141 |  | Y_R | -0.00052 |  | *_M | -3.65E-08 |
| M_N | -0.00140 |  | H_A | -0.00049 |  | W_Q | -2.98E-08 |
| S_H | -0.00131 |  | F_H | -0.00047 |  | *_T | -5.96E-11 |
| G_T | -0.00126 |  | W_A | -0.00040 |  | N_* | -5.99E-13 |
| K_A | -0.00121 |  | Q_I | -0.00030 |  | T_* | -4.14E-13 |
|  |  |  |  |  |  | *_D | -3.18E-13 |

**354 features with positive weight**

| **Table 2:** Features with positive weights. | | | | | | | |
| --- | --- | --- | --- | --- | --- | --- | --- |
| **Feature** | **Weight** |  | **Feature** | **Weight** |  | **Feature** | **Weight** |
| Consequence_SL | 4.23316 |  | Ref_A | 0.13240 |  | Consequence_DN | 0.06379 |
| SG_relCDSpos | 3.75932 |  | Consequence_U5 | 0.13043 |  | Consequence_IG | 0.06319 |
| SN_protPos | 0.86016 |  | I_GERPN_all | 0.12956 |  | F_Y | 0.06261 |
| protPos | 0.77082 |  | Ref_T | 0.12953 |  | Consequence_S | 0.06228 |
| oAA_E | 0.67582 |  | R_R | 0.12476 |  | UP_GERPN_all | 0.05978 |
| oAA_L | 0.63273 |  | Consequence_UP | 0.11148 |  | Q_K | 0.05674 |
| SG_relprotPos | 0.62898 |  | GERPN_all | 0.11024 |  | D_D | 0.05316 |
| NS_relprotPos | 0.38813 |  | A.1_T.1 | 0.10547 |  | G_E | 0.05314 |
| Consequence_NS | 0.34367 |  | T.1_A.1 | 0.10351 |  | Q_E | 0.05312 |
| Consequence_SN | 0.29906 |  | *_* | 0.10165 |  | U3_GERPN_all | 0.05103 |
| NS_protPos | 0.22639 |  | relprotPos | 0.10114 |  | NS_phylop_all | 0.04937 |
| IND_cDNApos | 0.20118 |  | Consequence_NC | 0.10050 |  | oAA_Q | 0.04901 |
| S_relCDSpos | 0.19874 |  | SG_PhyloP | 0.09185 |  | S_C | 0.04737 |
| S_protPos | 0.19605 |  | K_K | 0.09154 |  | Q_Q | 0.04650 |
| CS_relprotPos | 0.19470 |  | R_K | 0.08529 |  | oAA_K | 0.04639 |
| SN_relCDSpos | 0.17853 |  | C_C | 0.08323 |  | oAA_G | 0.04633 |
| Consequence_U3 | 0.16953 |  | Y_Y | 0.07912 |  | SN_PhyloP124 | 0.04596 |
| Consequence_I | 0.15362 |  | A.1_G.1 | 0.07284 |  | oAA_C | 0.04315 |
| SN_GERPRS_all | 0.15244 |  | T.1_C.1 | T.1_C.1 |  | SN_PhastCons | 0.04304 |
| relCDSpos | 0.14624 |  | T_S | T_S |  | NS_PhyloP124 | 0.04219 |

| **Feature** | **Weight** |  | **Feature** | **Weight** |  | **Feature** | **Weight** |
| --- | --- | --- | --- | --- | --- | --- | --- |
| A_S | 0.04175 |  | S_G | 0.02000 |  | NS_ReMap | 0.01047 |
| DN_GERPN_all | 0.04172 |  | G_W | 0.01985 |  | Y_C | 0.01023 |
| IG_GERPN_all | 0.03985 |  | P_Q | 0.01970 |  | Q_Y | 0.01018 |
| A_G | 0.03972 |  | CS_PhyloP124 | 0.01905 |  | P_K | 0.01009 |
| CS_GERPN_all | 0.03939 |  | S_N | 0.01886 |  | P_D | 0.01001 |
| oAA_Y | 0.03812 |  | I_V | 0.01849 |  | SG_ReMap | 0.00993 |
| RepeatType_LINE | 0.03787 |  | A_D | 0.01836 |  | H_K | 0.00974 |
| A_E | 0.03721 |  | D_E | 0.01813 |  | Y_M | 0.00967 |
| Grantham | 0.03560 |  | oAA_D | 0.01813 |  | S_D | 0.00956 |
| F_C | 0.03535 |  | W_L | 0.01806 |  | S_M | 0.00956 |
| M_L | 0.03485 |  | W_* | 0.01797 |  | H_G | 0.00908 |
| P_A | 0.03406 |  | K_Q | 0.01781 |  | oAA_P | 0.00899 |
| G_G | 0.03362 |  | SL_PhastCons | 0.01762 |  | V_N | 0.00899 |
| oAA_F | 0.03339 |  | NC_GERPN_all | 0.01712 |  | W_S | 0.00895 |
| V_L | 0.03336 |  | H_H | 0.01698 |  | A_P | 0.00894 |
| I_L | 0.03331 |  | H_R | 0.01694 |  | T_V | 0.00859 |
| H_Y | 0.03329 |  | S_GERPRS_all | 0.01652 |  | oAA_N | 0.00848 |
| G_C | 0.03148 |  | T_K | 0.01626 |  | W_M | 0.00844 |
| oAA_* | 0.03111 |  | K_E | 0.01610 |  | C_P | 0.00835 |
| T_N | 0.03049 |  | G_K | 0.01589 |  | SN_PhastCons124 | 0.00817 |
| G_A | 0.03039 |  | SL_CDSpos | 0.01587 |  | G_F | 0.00800 |
| T.1_G.1 | 0.02875 |  | S_Y | 0.01565 |  | M_S | 0.00798 |
| U5_GERPN_all | 0.02834 |  | SG_PhastCons124 | 0.01564 |  | EP | 0.00772 |
| A.1_C.1 | 0.02821 |  | oAA_M | 0.01550 |  | G_D | 0.00771 |
| Repeats | 0.02792 |  | P_L | 0.01536 |  | G_V | 0.00765 |
| oAA_H | 0.02785 |  | Q_L | 0.01450 |  | D_G | 0.00763 |
| Y_F | 0.02754 |  | V_Y | 0.01439 |  | A_K | 0.00760 |
| K_R | 0.02732 |  | SL_protPos | 0.01411 |  | S_A | 0.00740 |
| SL_relcDNApos | 0.02716 |  | C_S | 0.01408 |  | H_F | 0.00738 |
| W_C | 0.02605 |  | V_V | 0.01408 |  | isTv | 0.00736 |
| N_S | 0.02546 |  | V_E | 0.01341 |  | W_D | 0.00727 |
| F_F | 0.02488 |  | V_I | 0.01321 |  | ingene | 0.00724 |
| NS_relcDNApos | 0.02470 |  | oAA_R | 0.01304 |  | I_K | 0.00724 |
| DN_phylop_all | 0.02408 |  | N_N | 0.01274 |  | Q_F | 0.00723 |
| H_N | 0.02325 |  | R_T | 0.01253 |  | Q_V | 0.00716 |
| IG_phylop_all | 0.02320 |  | CS_protPos | 0.01236 |  | G_S | 0.00714 |
| R_D | 0.02254 |  | H_D | 0.01230 |  | IG_ReMap | 0.00706 |
| R_M | 0.02176 |  | S_T | 0.01220 |  | M_D | 0.00700 |
| SL_PhyloP124 | 0.02169 |  | N_H | 0.01207 |  | oAA_S | 0.00695 |
| oAA_V | 0.02102 |  | H_L | 0.01204 |  | V_K | 0.00691 |
| P_T | 0.02101 |  | RepeatType_DNA | 0.01189 |  | C_M | 0.00687 |
| P_S | 0.02078 |  | S_S | 0.01166 |  | W_E | 0.00684 |
| Alt_C | 0.02066 |  | W_I | 0.01103 |  | CS_phylop_all | 0.00656 |
| oAA_W | 0.02050 |  | relcDNApos | 0.01099 |  | M_Y | 0.00654 |
| Alt_G | 0.02033 |  | NC_cDNApos | 0.01092 |  | RepeatType_LTR | 0.00638 |
| P_H | 0.02011 |  | P_V | 0.01083 |  | I_A | 0.00634 |

| **Feature** | **Weight** |  | **Feature** | **Weight** |  | **Feature** | **Weight** |
| --- | --- | --- | --- | --- | --- | --- | --- |
| N_D | 0.00618 |  | NC_ReMap | 0.00339 |  | G_L | 0.00132 |
| MGW | 0.00612 |  | E_I | 0.00333 |  | Domain_ndomain | 0.00127 |
| HelT | 0.00607 |  | UP_phylop_all | 0.00321 |  | M_C | 0.00123 |
| T_I | 0.00606 |  | T_Y | 0.00320 |  | A_H | 0.00119 |
| W_T | 0.00599 |  | RepeatType_No_repeat | 0.00314 |  | Q_N | 0.00116 |
| SL_GERPN_all | 0.00595 |  | C_V | 0.00308 |  | H_M | 0.00113 |
| T_A | 0.00578 |  | S_relcDNApos | 0.00304 |  | R_A | 0.00107 |
| DN_PhastCons124 | 0.00576 |  | U5_ReMap | 0.00303 |  | D_A | 0.00106 |
| S_PhastCons124 | 0.00557 |  | C_G | 0.00302 |  | A_F | 0.00101 |
| C_D | 0.00555 |  | *_C | 0.00291 |  | P_E | 0.00101 |
| A_N | 0.00553 |  | *_W | 0.00289 |  | C_Q | 0.00100 |
| F_K | 0.00542 |  | S_Q | 0.00281 |  | C_A | 0.00100 |
| W_Y | 0.00539 |  | P_G | 0.00269 |  | N_C | 0.00095 |
| N_T | 0.00534 |  | I_E | 0.00267 |  | RepeatType_Satellite | 0.00088 |
| R_E | 0.00533 |  | A_R | 0.00267 |  | P_N | 0.00086 |
| NC_PhastCons | 0.00531 |  | U3_cDNApos | 0.00265 |  | V_R | 0.00085 |
| C_Y | 0.00527 |  | I_C | 0.00263 |  | D_K | 0.00082 |
| L_E | 0.00525 |  | ProT | 0.00249 |  | N_Q | 0.00081 |
| C_L | 0.00514 |  | S_V | 0.00248 |  | H_I | 0.00078 |
| Q_D | 0.00505 |  | Y_L | 0.00237 |  | A_C | 0.00078 |
| W_F | 0.00505 |  | V_W | 0.00234 |  | V_C | 0.00077 |
| D_Q | 0.00496 |  | S_E | 0.00232 |  | P_I | 0.00076 |
| R_F | 0.00490 |  | N_Y | 0.00223 |  | G_I | 0.00075 |
| F_E | 0.00487 |  | P_F | 0.00220 |  | UP_ReMap | 0.00071 |
| F_D | 0.00458 |  | miRNA | 0.00214 |  | S_PhastCons | 0.00069 |
| CS_relcDNApos | 0.00450 |  | R_V | 0.00206 |  | SL_ReMap | 0.00067 |
| E_T | 0.00450 |  | T_C | 0.00204 |  | P_M | 0.00067 |
| *_E | 0.00445 |  | Consequence_CS | 0.00197 |  | V_P | 0.00065 |
| L_N | 0.00437 |  | DN_PhastCons | 0.00196 |  | T_H | 0.00065 |
| T_D | 0.00428 |  | T_L | 0.00189 |  | I_* | 0.00064 |
| U5_relcDNApos | 0.00427 |  | I_H | 0.00188 |  | UP_PhastCons124 | 0.00063 |
| S_K | 0.00427 |  | UP_PhastCons | 0.00180 |  | I_D | 0.00062 |
| M_I | 0.00422 |  | M_V | 0.00179 |  | H_S | 0.00054 |
| SN_cDNApos | 0.00418 |  | SL_GERPRS_all | 0.00178 |  | K_P | 0.00053 |
| C_N | 0.00412 |  | Q_P | 0.00175 |  | N_V | 0.00051 |
| *_P | 0.00410 |  | I_Y | 0.00174 |  | H_P | 0.00050 |
| A_L | 0.00403 |  | IG_PhastCons | 0.00164 |  | D_S | 0.00049 |
| S_cDNApos | 0.00402 |  | F_L | 0.00161 |  | H_E | 0.00049 |
| P_Y | 0.00402 |  | A_W | 0.00158 |  | N_L | 0.00039 |
| RepeatType_RC | 0.00396 |  | Q_G | 0.00149 |  | N_E | 0.00036 |
| Domain_UD | 0.00389 |  | P_C | 0.00142 |  | C_E | 0.00030 |
| T_E | 0.00362 |  | R_Y | 0.00142 |  | D_T | 0.00030 |
| E_W | 0.00360 |  | BG3_state | 0.00141 |  | M_Q | 0.00030 |
| M_K | 0.00350 |  | F_G | 0.00138 |  | N_R | 0.00029 |
| ReMap | 0.00347 |  | *_G | 0.00136 |  | Y_A | 0.00029 |
| E_H | 0.00343 |  | D_L | 0.00132 |  | A_Q | 0.00026 |

| **Feature** | **Weight** |  | **Feature** | **Weight** |  | **Feature** | **Weight** |
| --- | --- | --- | --- | --- | --- | --- | --- |
| U3_PhastCons | 0.00020 |  | D_M | 7.38E-05 |  | V_T | 2.13E-08 |
| M_G | 0.00016 |  | TFBS | 5.87E-05 |  | D_* | 1.56E-12 |
| H_T | 0.00015 |  | A_* | 5.78E-05 |  | E_M | 4.80E-13 |
| A_V | 0.00013 |  | N_A | 1.31E-05 |  | *_V | 4.47E-13 |
| T_G | 9.81E-05 |  | T_Q | 1.30E-05 |  | *_A | 3.90E-13 |
| H_* | 9.39E-05 |  | M_H | 2.13E-08 |  | *_N | 3.60E-13 |
